# Theoretical Quantification of the Polyvalent Binding of Nanoparticles Coated with Peptide-MHC to TCR-Nanoclusters

**DOI:** 10.1101/2022.03.04.483017

**Authors:** Manuela Pineros-Rodriguez, Louis Richez, Anmar Khadra

**Affiliations:** Department of Mathematics and Statistics, McGill University, Montreal, Canada; Quantitative Life Sciences Program, McGill University, Montreal, Canada; Department of Physiology, McGill University, Montreal, Canada

**Author notes:** Contributed equally to this work (Manuela Pineros-Rodriguez, Louis Richez). **Corresponding author:** Anmar Khadra, McIntyre Medical Building (room 1120), 3655 Prom. Sir William Osler, Montreal, Quebec, Canada H3G 1Y6.

**Keywords:** Polyvalent biding, pMHC-coated nanoparticles, T cell receptors, multiscale model, dissociation constant, Gibbs free energy, insertion probability, TCR nanoclusters, Density of bound and triggered TCRs, cooperativity

## Abstract

Nanoparticles (NPs) coated with pMHCs can reprogram a specific type of CD4+ T cells into disease-suppressing T regulatory type 1 cells by binding to their TCRs expressed as TCR-nanoclusters (TCR_nc_). NP size and number of pMHCs coated on them (called valence) can be adjusted to increase their efficacy. Here we explore how this polyvalent interaction is manifested and examine if it can facilitate T cell activation. This is done by developing a multiscale biophysical model that takes into account the complexity of this interaction. Using the model, we quantify pMHC insertion probabilities, dwell time of NP binding, TCR_nc_ carrying capacity, the distribution of covered and bound TCRs by NPs, and cooperativity in the binding of pMHCs within the contact area. Model fitting and parameter sweeping further reveal that moderate jumps between IFN*γ* dose-response curves at low valences can occur, suggesting that the geometry of NP binding can prime T cells for activation.

## 1 Introduction

T cells recognize and react to foreign antigens as well as, in the case of autoimmune diseases, to self-antigens through antigen-specific interactions between T cell receptors (TCRs) and peptide-major histocompatibility complexes (pMHCs) class I (in the case of CD8^+^ T cells) and class II (in the case of CD4^+^ T cells) [1, 2]. These pMHC molecules are typically expressed on the membrane of antigen presenting cells, such as dendritic cells and macrophages, for T cell activation, as well as on the surface of nucleated cells during both an infection or an autoimmune disease [1].

The binding of TCRs to pMHCs constitutes one key element of the immune synapse that forms between T cells and antigen presenting cells [3, 4]. Several quantitative studies have previously analyzed different kinetic aspects of TCR-pMHC interactions within the immune synapse, including pMHC mean residency time [5, 6] and TCR internalization [7]. The role of binding and unbinding rates in T cell activation through rebinding and confinement [8,9] together with antigen discrimination [10] have been also investigated quantitatively. These modeling studies typically incorporated kinetic proofreading [4,11] into their model formalisms when studying the sensitivity of TCRs to antigens. In this kinetic proofreading model, the continued TCR-pMHC engagement and subsequent phosphorylation of the TCR associated immunoreceptor tyrosine-based activation motifs (ITAMs) by signaling molecules [4] were taken into account [10]. The presence of these multiple modifications steps (through phosphorylations) allows TCRs to exhibit ultrasensitivity toward antigens [12]

It has been previously shown that administering nanoparticles (NPs) that are coated with disease-relevant cognate pMHC class II molecules are able to reprogram T helper type 1 CD4^+^ T cells (that are both endogenous and autoantigen-experienced) into disease-specific immunosuppressive T regulatory type 1 (Tr1) cells [13] and subsequently expand and recruit them to target organ and associated lymphoid tissue [14, 15]. These TR1 cells have autoregulatory and immunomodulatory effects that cause the suppression of inflammation in autoimmune diseases through complex immunological processes [16–22], providing evidence for its therapeutic potential as nanovaccines [15,23]. Using computational modeling approaches, the effects of such treatment on the progression of autoimmune diseases (e.g., type 1 diabetes) was also quantified, taking pMHC density on NPs along with NP concentration and frequency into consideration [23–26].

Upon binding to TCRs, pMHC-coated NPs aggregate on the surface of T cells [14]. Indeed, it has been demonstrated that, following stimulation, TCRs form tight arrangements, or micro-clusters, on T cell surface [27]. These micro-clusters can contain between 40 – 150 TCRs, and cover an area of 0.35 – 0.5 *μ*m [28–30]. Nonetheless, recent studies have shown that, even prior to stimulation, TCR are organized in smaller structures, or nanoclusters [30–32]. These contain up to 20 TCRs, distributed over regions of radius of up to 200 nm. While it remains unclear how these TCR nanoclusters (TCR_nc_) are formed, their presence appear to be essential for proper T cell activation [33]. Several hypotheses have been proposed on how this effect is manifested, including the induction of conformational changes on the TCR nanoclusters and cooperativity [32]. TCR-pMHC interactions thus take place within these TCR_nc_, and their known geometrical properties provide a framework that one can use to further decipher the polyvalent binding of pMHC-NPs with TCR_nc_.

The arrangement of TCR into TCR_nc_ complexes imposes constraints on the spatial distribution of TCR on the surface of T cells. A minimal distance thus must be preserved between TCRs, since the co-receptors associated with each of these TCRs occupy a region that cannot be occupied by any other. This minimal distance required between two TCRs is ~ 10 nm [34]. It has been shown that ligand-receptor binding can be significantly increased when either of these agents is presented in a polyvalent manner, through mechanisms like chemical cooperativity or mechanistic effects [35, 36]. In contrast, other studies have focused on the negative effect of spatial constraints in ligand-receptor binding [37–39], such as steric effects [40]; in these latter studies, it was determined how the dynamics on a ligand-receptor system are modified when the ligand is presented on an area of known geometry, and the receptor imposes structural constraints on the binding. Considering the effects of all these geometrical factors can help us understand the kinetics of NP binding to T cells, without the need to estimate any parameters.

Studying the effect of geometry on T cell activation - usually measured experimentally in terms of interferon-*γ* (IFN*γ*) release - can elucidate both how to optimize the design of NPs to maximize Tr1-cell response, and what the underlying dynamics of the TCR-pMHC interactions are. In this study, we perform such analysis using both modeling and computational techniques. To this end, we develop a multi-scale model of this system by incorporating the three levels of interactions between NPs and T cells to determine how geometry defines this polyvalent binding and cooperativity. By considering an affinity-centric model, which associates T cell activation to the interaction time between ligand (pMHC) and receptor (TCR), we explore if such a multiscale model can produce jumps between IFN*γ* dose-response curves generated by T cells upon NP stimulation when the valence of pMHC (total number of pMHCs on NPs), denoted by *v* is changed modestly for a given NP size. The goal is to determine if the geometry of interaction can facilitate T cell activation.

## 2 Methods

### 2.1 General Description of the Multiscale Model

To model pMHC-coated NPs binding to T cells at the population level, we employ a reverse engineering approach. This is accomplished by designing a multiscale model of T cell-NP interaction while ignoring TCR trafficking [41]. The model is comprised of three scales:

1. The *contact area model* (CAM) that describes the binding of a single NP of a specific size, coated with *v* pMHCs (referred to hereafter as valence), to one or more TCRs. By constraining TCR-pMHC interactions using geometrical considerations defined by the TCR_nc_ and the NP, and applying Monte Carlo simulations, we obtain distributions for the number of bound pMHCs per NP [40, 42]. These computations are detailed in the Results Section.
2. From the distributions determined by the CAM, a *TCR nanocluster model* (TNM) is developed to describe how multiple NPs bind to a TCR_nc_. The model should include the adsorption and desorption rates of NPs. The dynamics considered at this level yield steady-state solutions for the number of bound NPs per TCR_nc_ obtained by applying the continuation method (detailed in the Results Section). The results of the NC model can then be scaled to the entire cell to obtain the number of bound NPs per single T cell.
3. Finally, the steady states of the TNM are fed into a *T cell activation model* (TAM) that employs the kinetic proofreading model - associated with TCR phosphorylation (Fig. S1) - to characterize the proportion of bound TCRs involved in the activation of a T cell leading to the production of IFN*γ*.

Table S1 provides a list of all symbols used to describe each model scale mathematically.

### 2.2 Kinetic Proofreading and T Cell Activation

While TNM returns the number of NPs attached to the cell at steady state, CAM yields the number of TCRs bound to those NPs. It now remains to translate the steady state number of bound TCRs to an actual production of IFN*γ*. This is done through the use of TAM, in which the formation of a TCR-pMHC complex induces a series of phosphorylation steps to each TCR [11]; when these TCRs reach their final phosphorylation step, they get triggered, allowing them to contribute to T cell activation. Assuming the receptor must undergo a number, *κ*, of these phosphorylation steps, each with the same rate *k_p_*, then according to the kinetic proofreading model (Fig. S1), the likelihood of triggering is given by

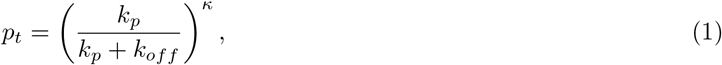

where *k_off_* is the off-rate of pMHC unbinding from a TCR, as opposed to the on-rate *k_on_* (both of which have units of time^-1^ and typically referred to as the cross-linking rates [37]). Once the number of triggered TCRs per T cell is determined, the actual quantity of IFN*γ* produced, denoted by *y*(*x*) (where *x* is the number of triggered TCRs), is calculated using the expression

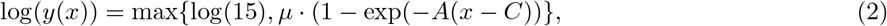

where *μ* is the maximum level of activation, *A* is the rate of activation increase and *C* is the activation threshold. The log(15) term accounts for the basal level of IFN*γ* release by a T cell. The expression in Eq. (2) is chosen to mimic the profile of IFN*γ* release from a T cell obtained upon stimulation with a peptide, antigen or CD3 as a function of triggered TCRs [43]. According to this profile, a threshold of triggered TCRs per T cell must be surpassed to induce activation; the threshold is followed by a steep increase in T cell activation until reaching a plateau (see Fig. S2). [For comparisons between the graph of Eq. (2) and IFN*γ*-release data, see Fig. 4.1 in [42].]

If 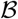 is the number of bound TCRs per T cell, then the number of triggered TCRs satisfies 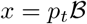. By Eq. 2, it follows that

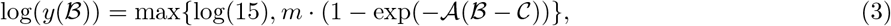

where 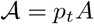 and 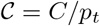.

### 2.3 Model Parameters and Numerical Computations

The resulting multiscale model possesses five parameters. These are: the dissociation constant *K_D_* = *k_off_*/*k_on_* of one single pMHC from a TCR, the dissociation constant 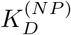 of one single NP from a TCR_nc_, the parameters *μ*, 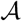 and 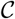 associated with the IFN*γ* release as a function of triggered TCRs as detailed in Eq. (3).

The values of these parameters are estimated using Metropolis-Hasting method in the MCMC toolbox (available online: https://mjlaine.github.io/mcmcstat/) for MATLAB (Mathworks, Natick, MA, USA). The doseresponse curves of IFN*γ* release, obtained from T cells stimulated with pMHC-coated NPs [23], are used to estimate model parameters, 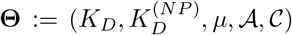, with the goal of minimizing the sum-of-squares error (SSE), given by

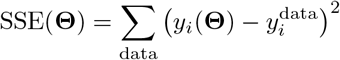

where 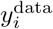 represents the level of IFN_*γ*_ associated with a single (*r*, *v*, *D*) triplet and *y_i_*(**Θ**) its corresponding model prediction. Fitting is performed separately on the *r* = 14 and *r* = 20 nm data for 15000 iterations of the MCMC algorithm. At each iteration *i* with parameter values **Θ**_*i*_, the Metropolis-Hasting method proposes a candidate set of parameter values 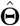 sampled from a normal distribution centered at **Θ**_*i*_. The candidate parameter set is accepted with probability 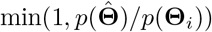, where

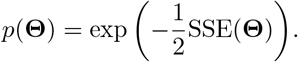

This method ensures that the chain will preferentially generate estimates lowering the SSE, yet while still exploring a wide range of the parameter space.

Details of this fitting are shown in Fig. S3 (for more details, see TAM outcomes in Results). The codes for regenerating the figures are available online [44].

### 2.4 Interferon-*γ* (IFN*γ*) Data

In order to explore how NP design (including radius and valence) affects T cell activation, the Santamaria Lab (University of Calgary) incubated 25000 CD8^+^ T cells for 48 h with NPs coated with cognate pMHCs at different concentrations, and with various radius (*r*) and valence (*v*) pairs. They then measured T cell response in terms of interferon-*γ* (IFN*γ*) secretion [23]. The NP-dependent dose-response curves for IFN*γ* release, obtained using NPs with two different radii (*r* = 14 and *r* = 20 nm), are displayed in Fig. S3. This figure shows that increasing the NP concentration gradually and steadily increases T cell response, and that the latter is enhanced upon increasing the valence. A nonlinear jump is observed between the two dose-response curves associated with valence *v* = 8 and *v* = 11 for *r* = 14 nm, which the Santamaria Lab hypothesized to be due to crossing an activation threshold of pMHC density.

## 3 Results

### 3.1 Outcomes of the Contact Area Model (CAM)

#### 3.1.1 CAM Design and Formalism

The CAM imposes most of the geometrical constraints of the polyvalent interaction between a NP and a clustered set of TCRs. In particular, when a NP first binds to a T cell, only a portion of the NP surface area becomes available for interaction with the T cell.

This region, known as the contact area, is defined by an angle *θ* representing how wide it is relative to the center of the interaction (Fig. 1). The larger the NP (i.e., the bigger the NP radius *r*), the larger the contact area. We choose *θ* to be 45°measured from the vertical axis. The consequence of this constraint is that only the pMHC contained within the contact area are available to bind to the TCRs, a quantity known as the effective valence 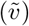. The choice of angle will be motivated when discussing how it influences the effective valence 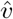. With *θ* = 45°, we have 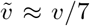 [23]. As stated before, changing NP radius will alter the effective valence 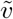 due to change in pMHC density on NPs.

**Figure 1:**
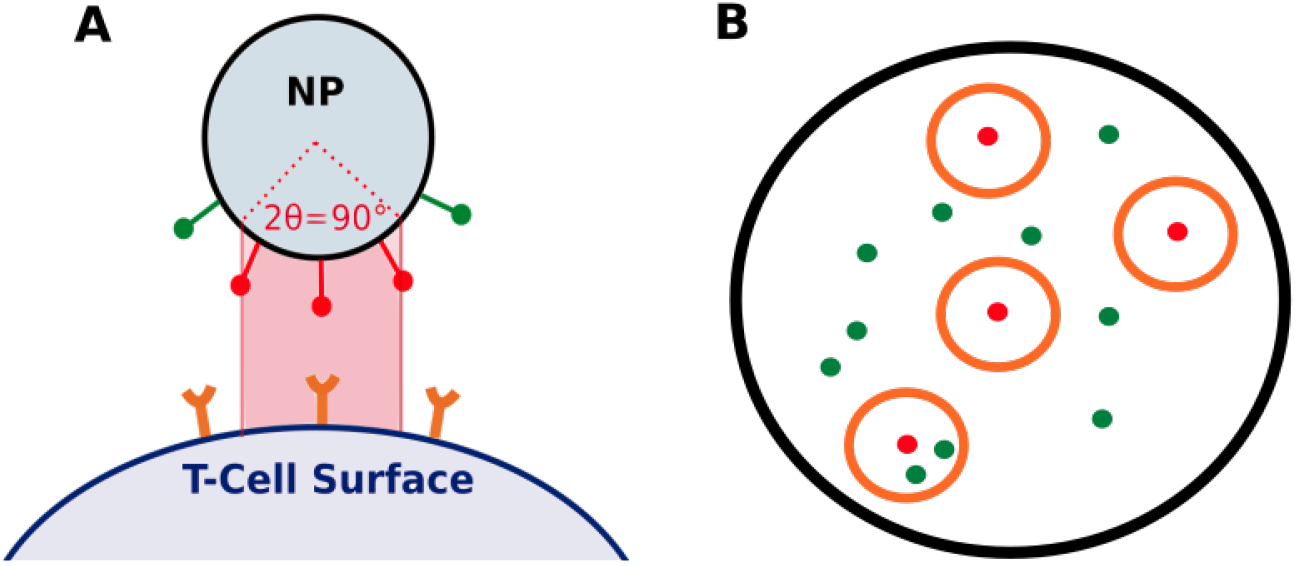
Schematic of the contact area. (A) The geometry of the contact area (red shaded region) constrains the number of pMHCs available for binding, a quantity called the effective valence 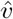 (depicted as red pMHCs). (B) A bird’s-eye view of the contact area showing a random configuration of TCRs (orange circles) and pMHCs (red dots: bound to TCRs; green dots: unbound).

The second set of geometric constraints to be considered has to do with the density of TCRs on T cell surface. TCR density is determined by three quantities in CAM: the minimum distance between TCRs (*d_TCR_* = 10 nm), the radius of TCR nanoclusters (*r_nc_* = 100 nm) and the number of TCRs per nanocluster (*n_TCR_* = 20) [32]. In this study, we will consider a range of possible TCR densities to investigate how it impacts CAM dynamics.

By letting

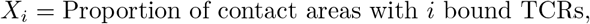

we can design a Markov model that describes the transitions between these states representing pMHC binding and unbinding events (Fig. 2). The dynamics of this model is given by

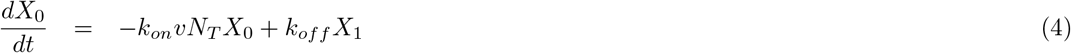

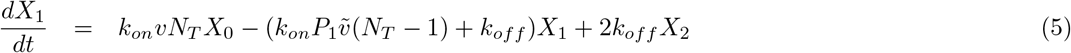

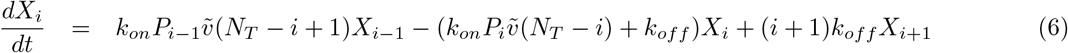

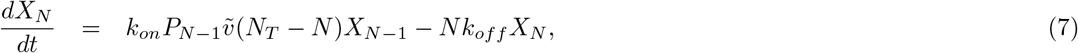

where *k_on_* and *k_off_* are the cross-linking rates defining the monovalent binding and unbinding rates of a single TCR-pMHC bond, *N_T_* is the number of TCRs in the contact area (referred to hereafter as the “number of covered TCRs” by a NP), and *P_i_*, *i* = 1,…, *N* – 1, is the insertion probabilities denoting the proportion of pMHCs that remain available for subsequent binding [40]. The geometrical constraints of CAM are encoded in the insertion probabilities *P_i_* (Fig. 1B), and they are estimated using Monte Carlo simulations of randomly generated configurations of pMHCs and TCRs within the contact area (as described in detail in the next section). If *N_P_* is the number of pMHCs in the contact area, then the maximum possible states (*N*) is given by

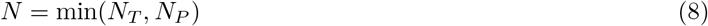

**Figure 2:**
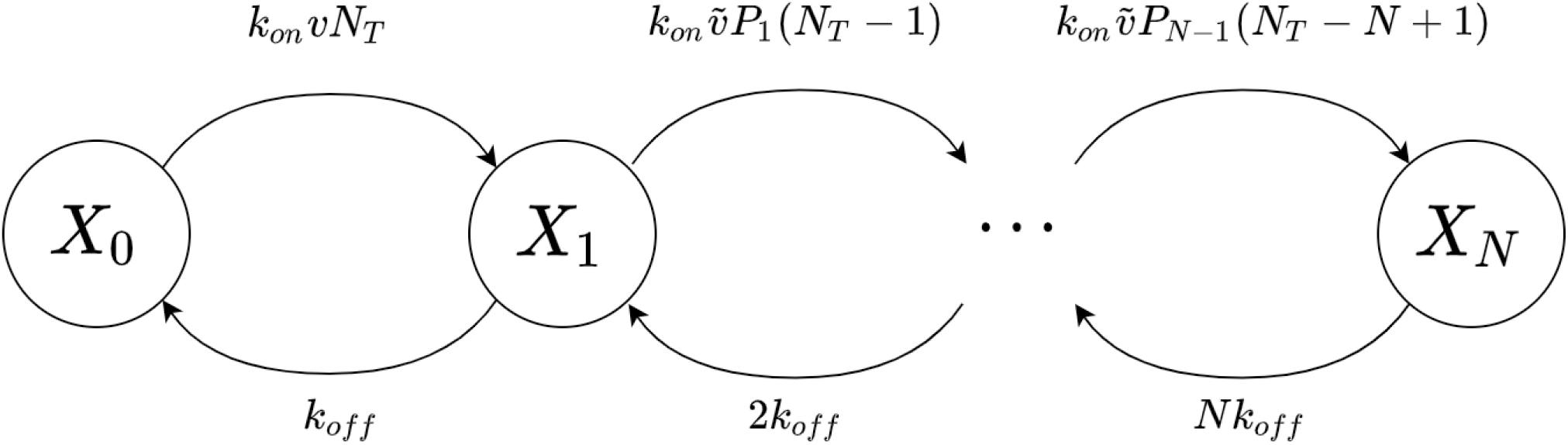
Markov scheme of the contact area model (CAM). Each state *X_i_*, *i* = 0,…, *N* represents the class of all contact areas with *i* bound pMHCs, where *N* is given by Eq. (8). Forward and backward transitions thus correspond, respectively, to single pMHC binding and unbinding events with rates as described by the expressions next to each arrow. Here, *P_i_* (*i* = 1,…, *N* – 1) denotes the insertion probabilities, *v* the NP-valence, 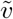 the effective valence, *N_T_* the number of TCRs in the contact area and *k_on_* and *k_off_* the cross-linking on and off rates, respectively.

For conciseness, we let *f_i_k_on_* and *b_i_k_off_* denote the forward and backward rates of the Markov model in Eqs. (4)–(7), where

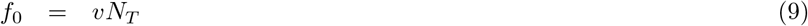

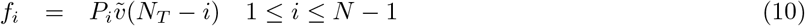

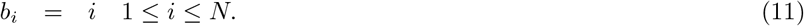

The average number of bound pMHC can be determined from the steady states of *X_i_*, denoted 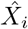. Setting Eq. (4) to zero, we obtain

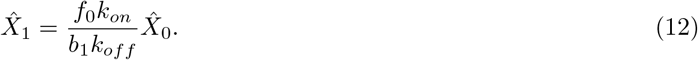

Substituting this expression into Eq. (5) and solving for the steady state yields

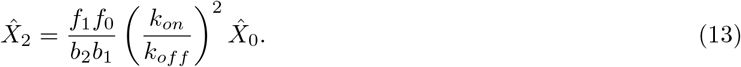

Continuing this process inductively through Eqs. (4)–(7) and setting

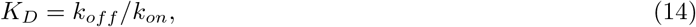

we obtain a general form for the steady state solutions of *X_i_*, given by

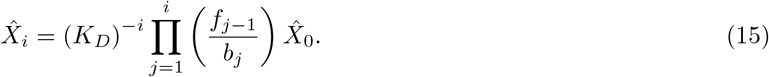

Since T cell half-life is on the order of days, a much longer time scale compared to that for NP binding to T cells, one can ignore T cell turnover in the model and assume that the system is conserved. Based on this, we have

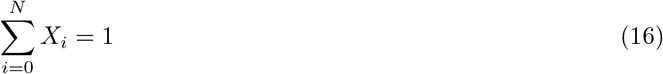

Combining this constraint with our result from Eq. (15), we can solve explicitly for 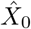 to obtain

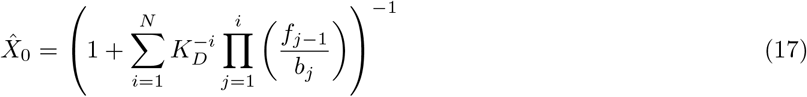

Plotting the steady states 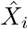 as a function of the dissociation constant *K_D_* = *k_off_*/*k_on_* for various combinations of NP radii (*r*) and valences (*v*) shows that for *r* = 14 nm (Fig. 3A), CAM ends up forming only two steady states: 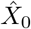 that increases with *K_D_*, and 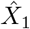 that decreases with *K_D_*, regardless of the valence *v*. Modifying the latter shifts the two steady state curves of 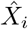, upward, for *i* = 1, and downward, for *i* = 0. The presence of only a single occupied steady state in this case is mainly due to the presence of a single TCR in the contact area. That is not the case for *r* = 20 nm, where a gradual increase in the valence from *v* = 9 to *v* = 210 increases the number of occupied steady states (Fig. 3B) due to the presence of multiple TCRs in the contact area. Interestingly, for higher valences, including *v* = 61 and *v* = 210, some steady states (e.g., the steady state 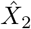 for *v* = 210), distinct from 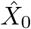, exhibit increasing profiles with *K_D_*. This seems to suggest that certain geometries are preferred for a given range of *K_D_* values and pairs of (*r*, *v*). For large values of *K_D_*, these monotonically increasing steady state curves eventually begin to decline, implying that such geometries become no longer preferred (results not shown).

**Figure 3:**
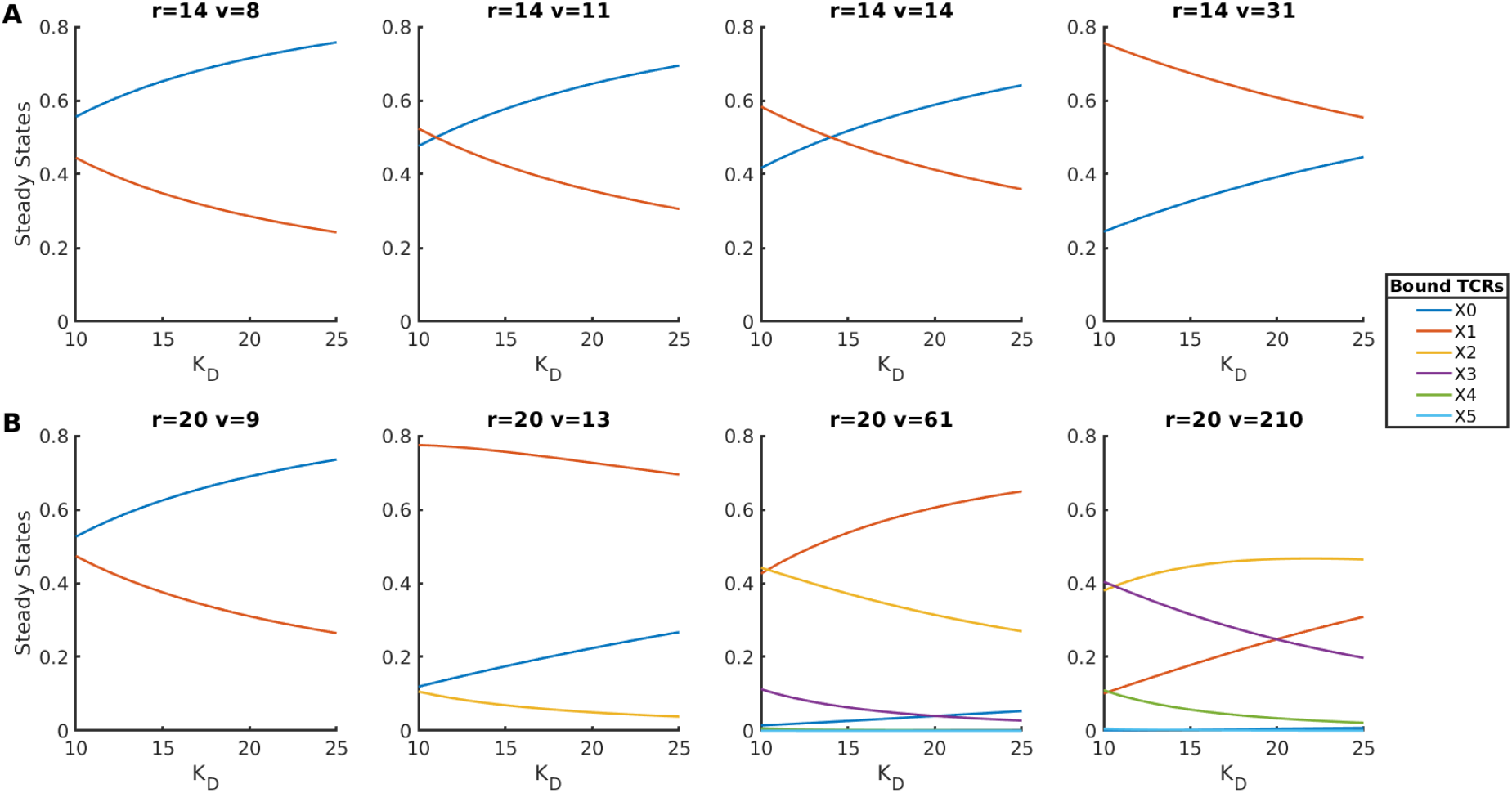
Steady states of the proportion of contact areas with *i* bound TCRs. Profiles of 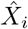, *i* = 1,…, *N* of CAM plotted against *K_D_* for (A) *r* = 14 nm (*v* = 8, 11, 14, 31), and (B) *r* = 20 nm (*v* = 9, 13, 61, 210) NPs. Curves are color-coded according to the legend. Notice how for *r* = 14 nm, NPs are limited to a single occupied steady state since only a single TCR is contained in the contact area.

#### 3.1.2 Monte Carlo Simulation for Computing Insertion Probabilities

The insertion probabilities *P_i_*, *i* = 1,…,*N*, embedded in the transition rates of the Markov Model in Eqs. (4)–(7), are estimated by Monte Carlo simulations of the contact area geometry (see Fig. S5). For a given configuration of pMHCs and bound TCRs, *P_i_* represents the proportion of remaining pMHCs available for subsequent binding [40].

They are computed as follows. We first generate a random configuration of 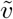 pMHCs within the contact area (Fig. S5I), followed by a random initial configuration of *i* TCRs (Fig. S5II) that are bound to specific pMHCs. TCRs are more likely to bind to regions with high pMHC density. Overlapping TCRs that could result from this step are re-positioned to ensure that the minimum separation between TCRs is satisfied (Fig. S5III). This could shift the TCRs to a low-density region of pMHCs.

To prioritize high density regions of pMHCs, we further define a neighborhood around each TCR possessing a radius *r_nbh_* = 2*d_TCR_* (Fig. S5IV). Other pMHCs within this neighborhood are considered “neighbors”. The locations of TCRs are then updated at each step to maximize the number of neighbors (Fig. S5V). The Monte Carlo loop allows the TCR to randomly relocate to one of the pMHC neighbors with a probability min(*N_*_/*N*_0_, 1), where *N*_0_* and *N*_*_ are the number of neighbors at the original and the new candidate site of binding, respectively. This ensures that the stationary distributions of TCRs on the contact area will correspond to regions with the highest density of pMHCs. Once the TCRs evolve to their stationary distributions, the insertion probabilities *P_i_* (*i* = 1,…, *N* – 1) are then calculated from the remaining pMHCs that are available for binding (Fig. S5VI).

By repeating these Monte Carlo steps for 5000 random configurations and averaging the mean proportion of available pMHCs over all trials, we obtain the insertion probabilities *P_i_*, where *i* is the number of bound TCRs within the contact area (Fig. 4). We do so for every (*r*, *v*) pair used in generating NP-dependent IFN*γ* dose-response curves (Fig. S3). Our results reveal that between *v* = 8 and *v* = 11, there is a prominent jump in the profiles of *P_i_* for *r* = 14 nm, but this prominent jump occurs between *v* = 9 and *v* = 13 for *r* = 20 nm. The increase in the effective valence 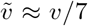 from a single to two pMHCs for both radii is the main reason we observe this jump in the insertion probabilities, highlighting the direct dependence of the latter on the contact area geometry. Furthermore, the profiles of *P_i_* (Fig. 4) indicate that when the pMHC valence increases, *P_i_* become increasingly dependent on the occupation of ligand sites. More specifically, each NP seems to reach a maximum number of simultaneously bound pMHCs, previously denoted by *N*. This quantity can be estimated by the maximum *i* at which the insertion probability *P_i_* > 0.01. This corresponds to the last state that has a probability larger than 1% of being transitioned to from the previous state. Reaching states with probability smaller than 1% is considered too unlikely to contribute to the dynamics of the system.

**Figure 4:**
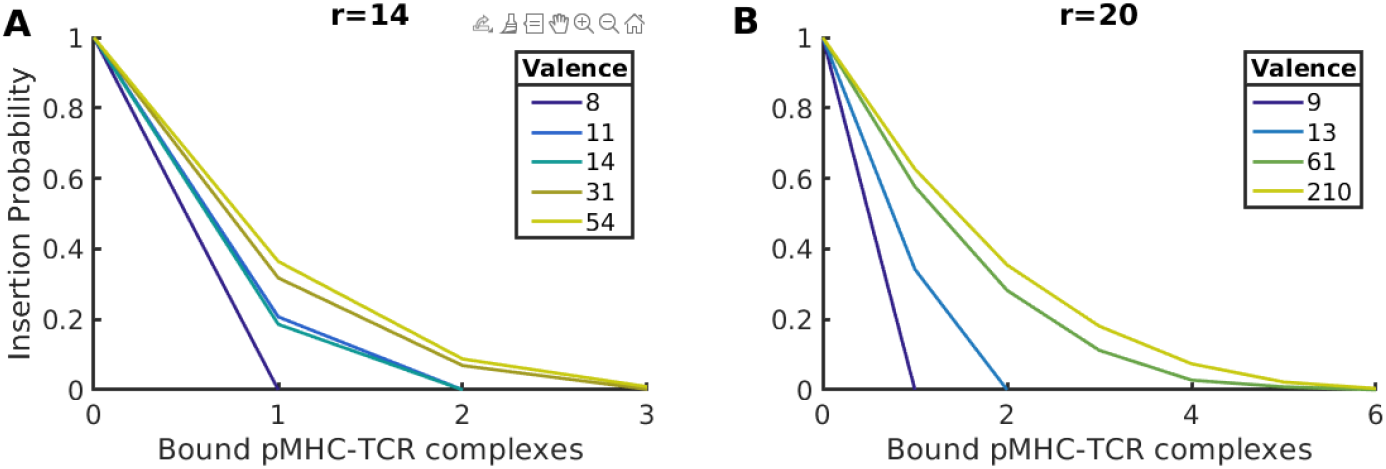
Insertion probabilities associated with CAM. The proportion of pMHCs that remain available for binding within the contact area after a given number of TCR-pMHC complexes are already formed are estimated from Monte Carlo sampling for (A) *r* = 14 nm, and (B) *r* = 20 nm NPs, and for all valences *v* listed in the legends. Curves are color-coded according to the legend in each panel.

#### 3.1.3 Quantifying NP Dwell Time

The transition rates of CAM also enable us to quantify analytically the dwell time of a NP on the contact area. i.e., the average time of interaction between a NP and a T cell. To do so, we first define

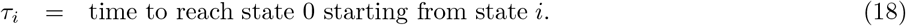

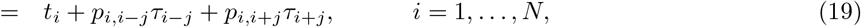

where *t_i_* is the average time spent in state *X_i_* and *p_i,j_* is the probability of transitioning from state *X_i_* to an adjacent state *X_j_* (only transitions to adjacent states are allowed). From model assumptions, we conclude that

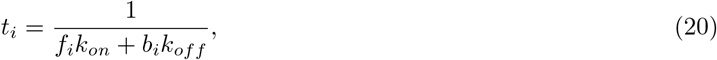

and that transition probabilities are given by

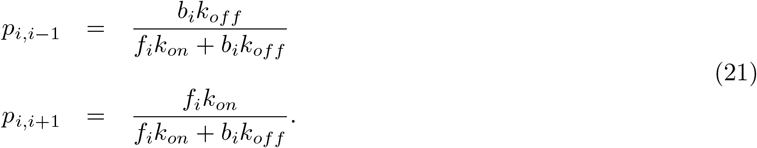

Substituting Eqs.(20) and (21) into Eq. (19), we obtain

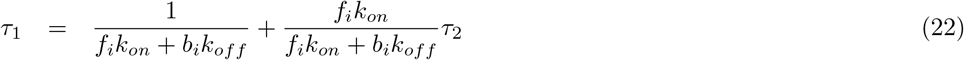

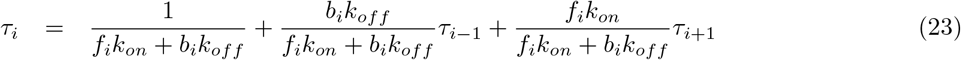

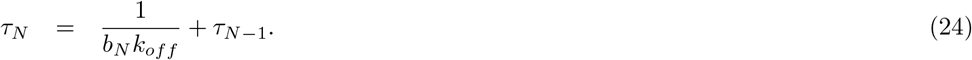

Rearranging the terms in Eqs. (22)–(24), we obtain this equivalent system of algebraic equations

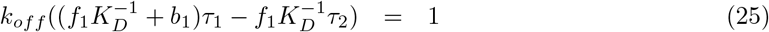

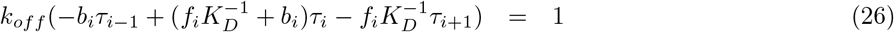

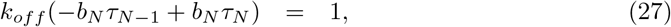

where as before *K_D_* = *k_off_*/*k_on_*. Rewriting Eqs. (25)–(27) in matrix form, we obtain

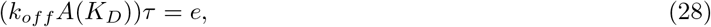

where *e* = (1,1, · 1)^*T*^ and

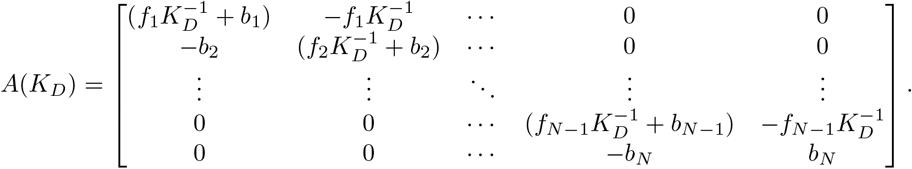

By solving for ***t*** = (*τ*_0_, *τ*_1_,…, *τ_N_*) using Eq. (28) and using steady states 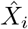, given by Eqs. (15) and (17), it becomes possible to compute the average dwell time 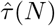 before a NP completely detaches from the TCR_*nc*_ (i.e. the time before all pMHCs on the NP become unbound). This can be done using the expression

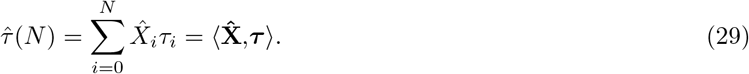

Numerically computing the average dwell times 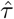 (Fig. 5) and plotting the results with respect to *K_D_*, for *r* = 14 nm (panel A) and *r* = 20 nm (panel B), generate, as expected, monotonically decreasing functions of *K_D_* for all valences *v*. Due to the fact that there is only one single TCR occupancy site in the contact area for *r* = 14 nm for all valences, the 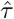 curves end up overlaying on top of each other in this case (Fig. 5A). In contrast, for *r* = 20 nm (Fig. 5B), the 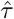 curves shift upward (especially at low *K_D_*) when increasing *v*, indicating more sensitivity to *v* due to the presence of multiple binding sites in the contact area. By artificially decreasing the minimal distance between two TCRs from its default value of *d_TCR_* = 10 nm to *d_TCR_* = 5 nm, allows the contact area formed by *r* = 14 nm NPs to have more than one occupancy site and thus higher number of steady states 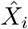 (Fig. S6); this causes the curves of average dwell time for various valences to produce outcomes similar to those seen with *r* = 20 nm. (Fig. S7B).

**Figure 5:**
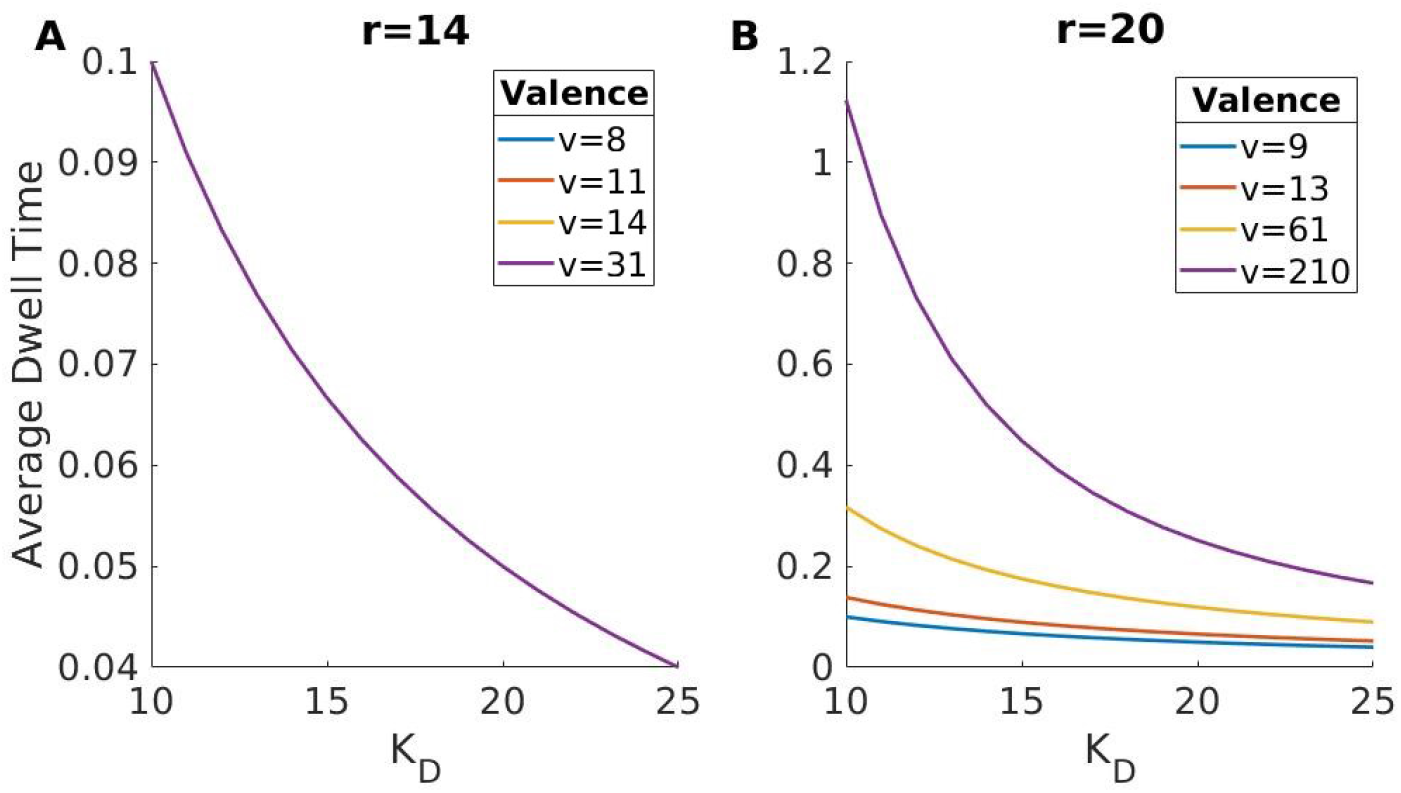
Dependence of average dwell time of a NP within the contact area on the dissociation constant and NP valence. Increasing the dissociation constant *K_D_* causes average dwell time 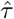 to monotonically decrease for both (A) *r* = 14 nm, and (B) *r* = 20 nm. However, increasing *v* causes 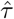 to exhibit no change to its profile when *r* = 14 nm (with all curves associated with valences listed in the legend lying on top of each other), or causes an overall elevation of its profile when *r* = 20 nm. Curves are color-coded according to the legend in each panel.

#### 3.1.4 Binding Cooperativity in CAM

The binding of a NP to a T cell is polyvalent in nature, governed by the interaction of single pMHCs with TCRs. This raises the question of whether pMHC binding to TCRs is cooperative at the contact area level, allowing single binding events to increase or decrease the likelihood of subsequent binding events. Generally speaking, Gibbs free energy provides a framework to compute cooperativity of a polyvalent interaction forming *N* complexes by expressing it in terms of the monovalent and polyvalent association constants *K^mono^* and *K^poly^*, respectively [35]. In the context of NPs and TCR_nc_, *K^mono^* corresponds to single interactions between pMHCs and TCRs, while *K^poly^* corresponds to overall interaction of all pMHCs and TCRs within the contact area as dictated by the cross-linking rates *k_on_* and *k_off_*. To compute cooperativity in the context of CAM, it is important to point out that this should be done in an already *established* contact area (adsorption and desorption of NPs are not involved). This means that the first forward transition *f*_0_ between *X*_0_ and *X*_1_ (Fig. 2) should depend on the effective valence 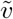, rather the *v*, i.e.,

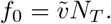

Biophysically, one can thus express cooperativity associated with a set of fixed *i* binding events, *i* = 1,…, *N*, within the contact area as follows

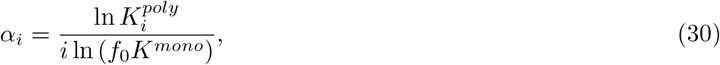

where *K^mono^* and 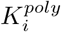 are given by

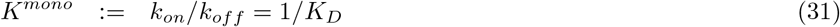

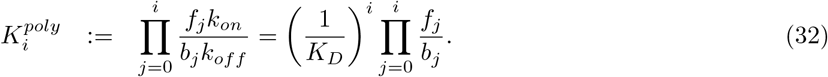

The *f*_0_ factor included with the monovalent association constant in Eq. 30 is necessary to account for the multiplicity of the first binding event proportional to 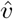 and *N_T_*. To consider all possible combinations of polyvalent binding that can occur within the contact area, the total cooperativity of a NP should be expressed as

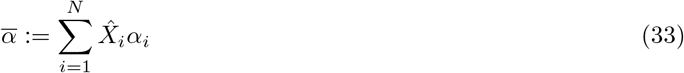

Here 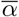 represents the average level of cooperativity that can exist between a single NP and its contact area. If 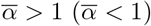, then cooperativity is positive (negative), with one single binding enhancing (impeding) the next. This handy rule of thumb, however, is valid only for values of *K_D_* < 1 (or *K^mono^* > 1).

Alternatively, one can characterize the type of cooperativity in CAM by simply considering the ratio of the kinetic rates between consecutive transitions in the model [45] using the equation

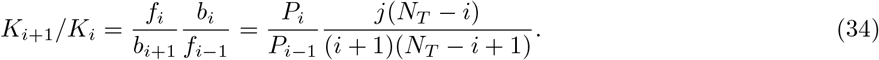

If each cross linking binding event is identical and independent (also known as statistical binding or neutral cooperativity) [45], then we expect the above ratio to satisfy

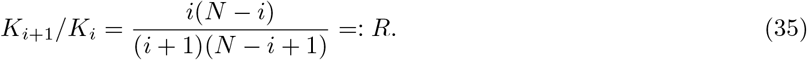

Equation (35) thus provides an alternative criterion for determining whether cooperativity is negative: *K*_*i*+1_/*K*_*i*_ < *R*, or positive: *K*_*i*+1_/*K_i_* > *R*. The merit of this criterion is that it provides a qualitative measure of cooperativity independent of the kinetic variable *K_D_*. Indeed, the crucial quantities in this measure are the insertion probabilities *P_i_*, *i* = 1,…, *N* – 1 that encode the geometric constraints of CAM.

There are two possible scenarios that one needs to consider when considering Eq. (35). This includes:

- **Case 1:** *N* = *N_T_* According to this scenario, TCR availability is the limiting factor in the polyvalent interaction within the contact area (i.e. 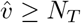), implying that the condition for negative cooperativity must simplify to

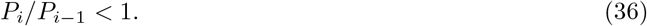 This condition is always satisfied since the insertion probabilities *P_i_* are monotonically decreasing.
- **Case 2:** 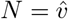 In this scenario, the number of pMHCs on NPs is the limiting factor (i.e. 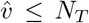). Accordingly, the condition for negative cooperativity becomes

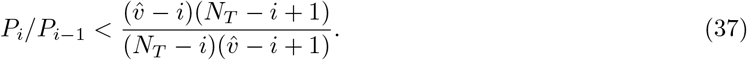

The above analysis allows us to determine the type of cooperativity that exists between pMHC binding events in CAM in two different ways. To compare the two approaches and assess cooperativity, we will evaluate 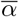 and link the results to the two cases obtained above. This is done first by computing 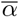 using Eq. (33) as a function of *v* = 10,…, 50, when *K^mono^* = 10, 20, 50, 100 (i.e., *K_D_* < 1). Plotting the results, (Fig. 6) reveals that there is an overall negative cooperativity in the binding of pMHCs to TCRs (*α* < 1) when *r* = 20 nm. In this case, cooperativity initially decreases with valences corresponding to conditions associated with the aforementioned Case 2 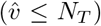 until eventually transitioning to Case 1. The valence at which this transition occurs depends on *N_T_* and therefore on the order of binding of the NP. Interestingly, despite the fact that there is an increase in cooperativity as *v* increases during Case 1, at no point does 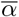 exceed neutrality 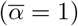 and enter the regime of positive cooperativity.

**Figure 6:**
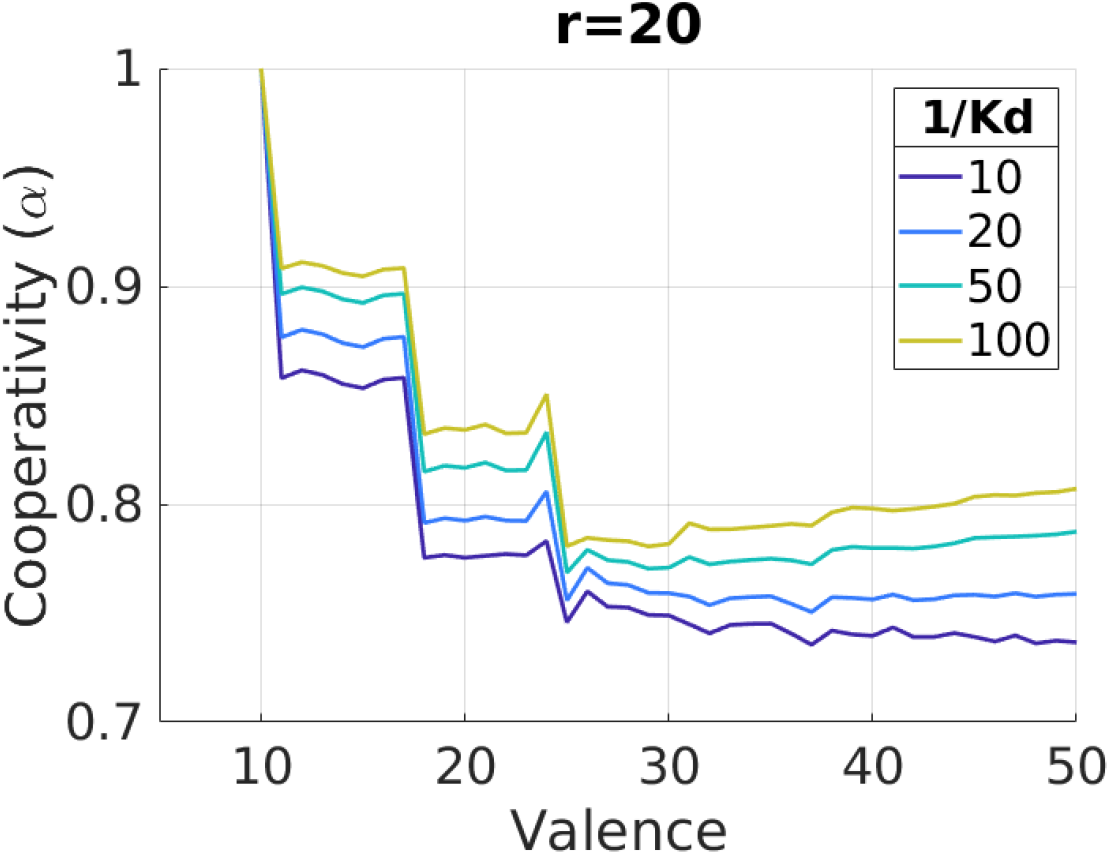
The average binding cooperativity defined by CAM. Average cooperativity 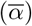 computed for *d_TCR_* = 10 nm and plotted as a function of NP-valence for various values of *K_D_* when *r* = 20 nm.

Because *d_TCR_* = 10 nm, there is only one single pMHC binding per *r* = 14 nm NPs, implying that there is no cooperativity in the binding of pMHCs within the contact area (results not shown). Setting *d_TCR_* = 5 nm, however, allows not only 20 nm NPs to exhibit negative cooperativity in pMHC binding, but also 14 nm NPs, with 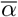 never exceeding neutrality to enter the regime of positive cooperativity (Fig. S8). The switching from Case 2 to Case 1 also occurs for both radii at *v* = 28 and *v* = 35 when *r* = 14 nm and *r* = 20 nm, respectively (Fig. S8A and B).

Taken together, these results imply that from a purely geometrical consideration, previous TCR-pMHC binding events do not promote the binding of subsequent TCR-pMHC complexes, and that the decrease in the availability of binding sites after each binding event is the main determinant of negative cooperativity.

#### 3.1.5 Optimizing the Angle of CAM

To justify the choice of the 45°angle chosen for CAM, we now investigate how altering this angle can affect the effective valence. We vary the angle *θ* of the contact area between 10°and 90°for all NP valences (Fig. 7). Our results reveal that, for relatively small but biophysically reasonable angles, the largest increase in effective valence between *v* = 8 and *v* = 11 for *r* = 14 nm NPs occurs at *θ* = 45° (Fig. 7A). The same result is obtained for *r* = 20 nm NPs (Fig. 7B). We can thus conclude that the 45° angle of the contact area is optimized and that changing the angle will not improve the outcomes of the model at higher levels.

**Figure 7:**
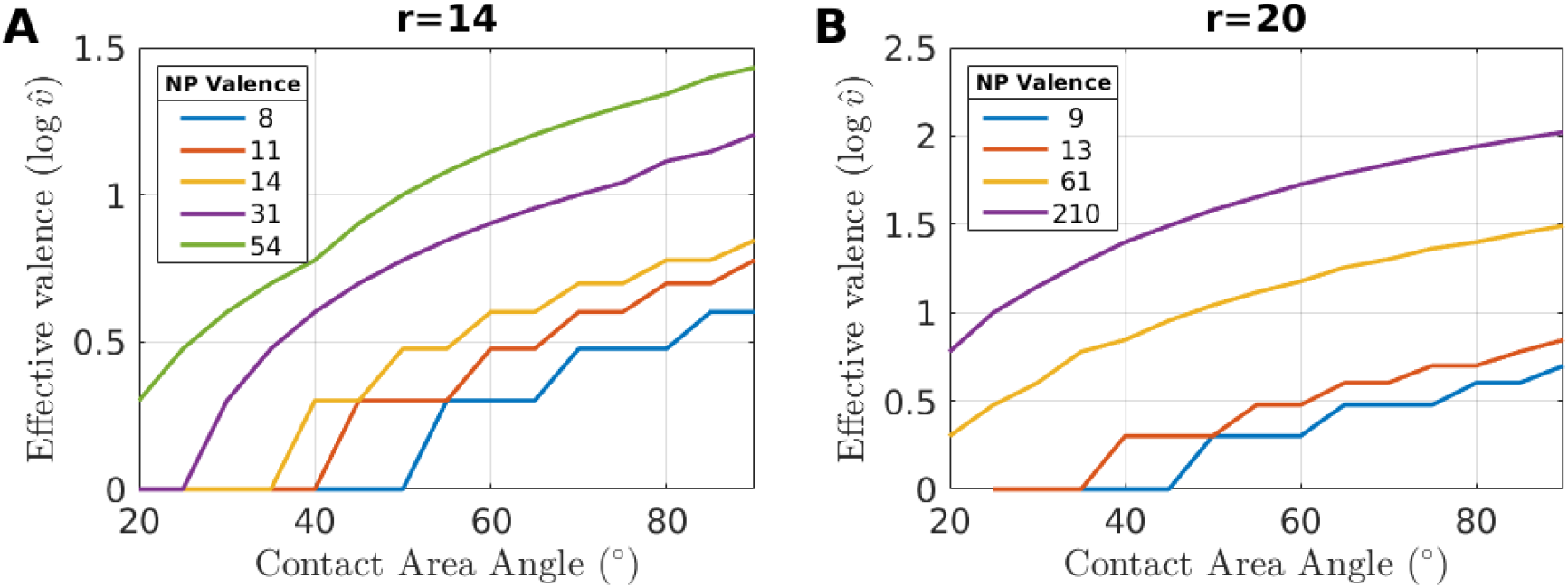
Dependence of effective valence on the contact area angle. Effective valence 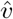 (in log scale) as a function the angle of the contact area *θ* for (A) *r* = 14, and (B) *r* = 20 nm NPs. Curves are color-coded according to the valences listed in the legend of each panel. Notice that when *θ* = 45°, there is a jump in 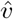 for both radii.

### 3.2 Outcomes of the TCR nanocluster model (TNM)

#### 3.2.1 TNM Design and formalism

TCRs are known to form nanoclusters on the surface of T cells (labeled TCR_nc_). By letting *M* denote the maximum number of NPs that could potentially bind to any TCR_nc_ configuration (i.e., the maximum carrying capacity), and by defining

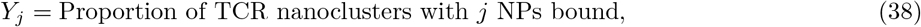

where *j* = 1,…, *M*, we can design a multi-state model that describes the dynamics of NP binding and unbinding to individual TCR_nc_ (Fig. 8). The transition rates between states in this model are the polyvalent binding, 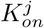, and unbinding, 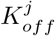 rates of NPs (*i* = 1,…,*M*). These forward and backward rates, respectively, are dependent on, but not necessarily equal to, the binding *k_on_* and unbinding *k_off_* rates of pMHCs to TCRs (due to the polyvalent nature of NP-TCR_nc_ interactions).

**Figure 8:**
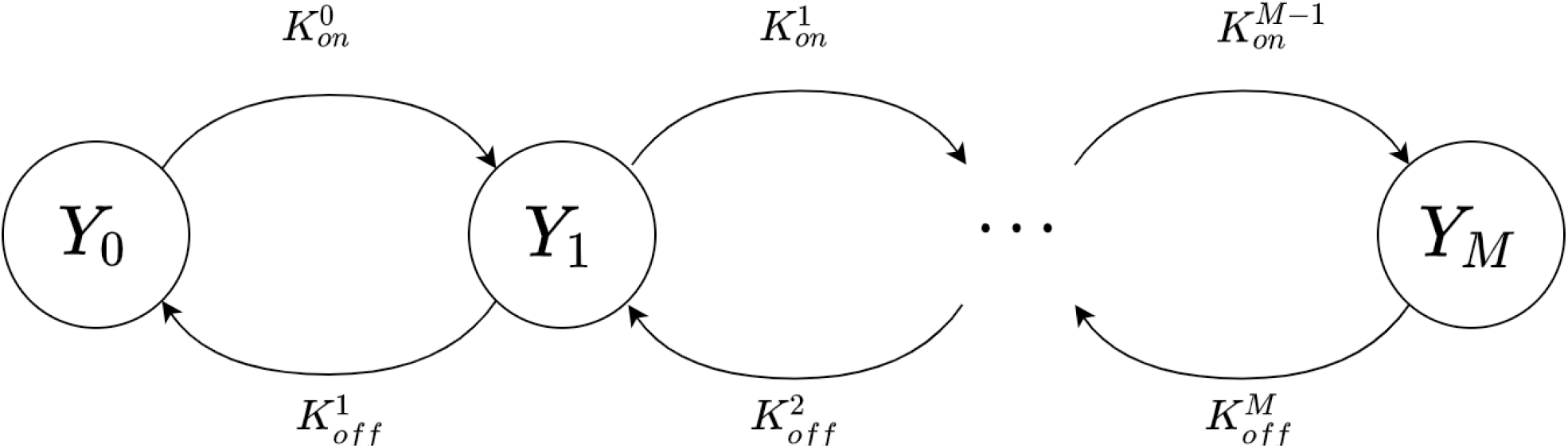
Multi-state scheme of the TCR nanocluster model (TNM). Each state *Y_j_*, *j* = 0,…, *M* represents the class of all TCR_nc_ with *j* bound NPs, where *M* is the maximum carrying capacity. Forward and backward transitions thus correspond, respectively, to single NP binding and unbinding events with rates 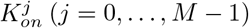 and 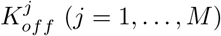.

The backward rates 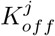 represents the rate at which one single NP detaches completely from a T cell; it depends on the inverse of the average dwell time 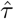 of the NP in the TCR_nc_ (given by Eq. (29)), which in turn depends on the number of covered TCRs *N_T_* by the NP. According to Fig. S5, *N_T_* depends on not only the position where the NP is bound on the TCR_nc_, but also on when the NP itself binds (first, second, etc.), implying that *N_T_* is a random variable. Letting

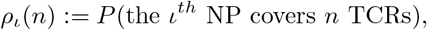

where *P* denotes probability, it follows that the average dwell time for the *ι^th^* NP bound to the TCR_nc_ is given by

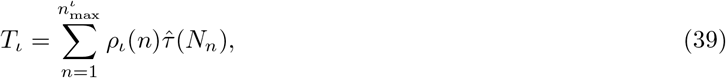

where 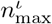 is the maximum possible number of simultaneously covered TCRs by the *ι^th^* NP and *N_n_* = min(*n*, *N_P_*) (see Eq. (8)). Based on this, we conclude that the unbinding rate of the *ι^th^* NP is 1/*T_ι_*. Given that the backward rate 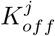 represents the unbinding of the first of *j* bound NPs to a TCR_nc_, we deduce that

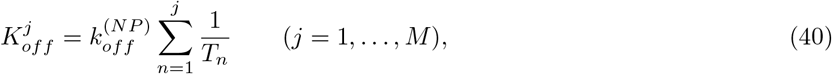

where 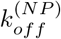 is the desorption rate of bound NPs.

The forward rates 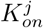, on the other hand, depend on the carrying capacity *c_NP_* for a given TCR_nc_ configuration, where *M* = max{*c_NP_*} evaluated over all possible configurations. The carrying capacity of a TCR_nc_ is simply the largest number of NPs that can be simultaneously bound to that given TCR_nc_. The *c_NP_* is dependent on the specific TCR configuration within the TCR_nc_, since denser TCR regions results in fewer bound NPs overall. We therefore expect the transition rates to states with higher indices to decrease and transitions to states with indices higher than *c_NP_* not to occur. We also expect the proportion of TCR_nc_ with a given *c_NP_* to be fixed. Let

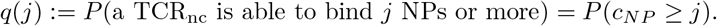

This quantity can be estimated from the Monte Carlo simulations of TCR distributions on a TCR_nc_. As the proportion of TCR_nc_ with *j* bound NPs (*Y_j_*) approaches *q_j_*, the binding of NPs becomes less likely. This behaviour should be represented by a saturation term in the TNM [42]. If the total number of TCR_nc_ per T cell is denoted by *m*, it follows that there are *mq_j_* TCR_nc_ per cell with a binding capacity of at least *j*. Since the binding rate of the NP depends on the NP concentration *D*, the valence of each NP *v* and the TCR-pMHC binding rate *k_on_*, we can deduce that the transition rate from state *Y_j_* to state *Y*_*j*+1_ is given by

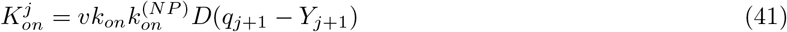

for *j* = 0,…, *M* – 1, where 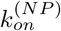 is the adsorption rate of NPs to a TCR_nc_, *q*_*j*+1_ – *Y*_*j*+1_ represents the saturation term we previously alluded to, and

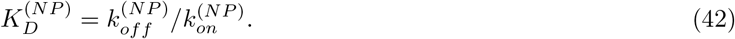

The mathematical framework of the multi-state model describing the dynamics of the TNM (Fig. 8) is thus given by

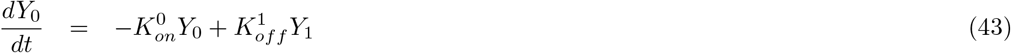

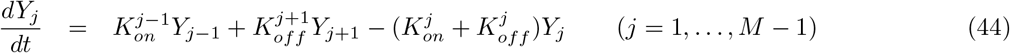

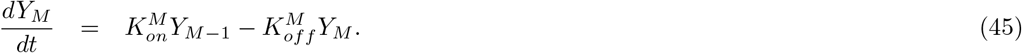

#### 3.2.2 Insertion Probabilities and Distributions of Covered TCRs According to TNM

Similar to CAM, TNM can be also used to compute the insertion probabilities associated with NP binding to a TCR_nc_. Using the same method outlined in Section 3.1.2, for a given number *i* of NPs bound to a TCR_nc_, we estimate via Monte Carlo simulations the proportion 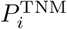 of remaining TCRs on the TCR_nc_ that are available for binding subsequent NPs averaged over 50,000 TCR_nc_.

The insertion probabilities for the TNM additionally provides estimates for *M*, the theoretical maximum carrying capacity of a TCR_nc_. This in turn serves as an input for another Monte Carlo simulation to directly estimate *q*(*j*) and the distribution of TCRs covered by a NP, *ρ_ι_*. Running this simulation for 50,000 possible TCR_nc_ configurations yields distributions of covered TCRs based on the order of binding of each NP. Recall that the first NPs will preferentially bind to regions with higher TCR densities, with subsequent NPs consequently covering fewer TCRs on average. These *ρ_ι_* can thus be determined using the expression

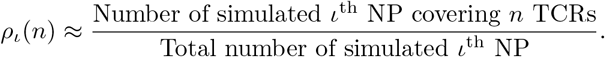

Furthermore, these simulations provide us with a probability estimate for the ability of a TCR_nc_ to bind at least *j* NPs or more, calculated as follows

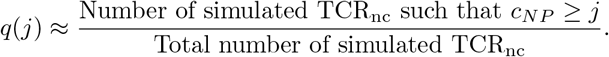

As outlined in the previous section, these quantities are necessary for defining the forward (*K_on_*) and backward (*K_off_*) rates of TNM and serve to encode the geometrical constraints associated with TCR_nc_-level dynamics.

In these computations, *M* is numerically calculated by quantifying the maximum *i* for which 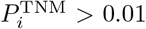. Beyond this *i* = *M*, the proportion of remaining TCRs is considered too unlikely to bind to any NPs. Plotting these insertion probabilities as a function of the number of bound NPs to TCR_nc_ (Fig. 9, blue curves) for both *r* = 14 (panel A) and 20 nm (panel B) NPs, we find that the maximum carrying capacity for *r* = 14 nm NPs is higher than that for *r* = 20 nm NPs. This is to be expected in view of the fact that *r* = 20 nm NPs are larger, requiring fewer NPs to cover the TCR_nc_ relative to the *r* = 14 nm NPs. Interesting, plotting the proportion of simulated TCR_nc_ that are able to bind *i* NPs for both radii (Fig. 9, orange curves), determined from the same Monte Carlo simulations used to compute *ρ_ι_*(*n*) and *q*(*j*), reveals that most, if not all, TCR_nc_ are able to bind at least several NPs. In fact, for both *r* = 14 (panel A) and *r* = 20 nm (panel B) NPs, most TCR_nc_ have carrying capacities close to the theoretical limit. This is deduced from the sharp drop in the orange curves as the number of bound NPs approaches *M* (dashed vertical lines).

**Figure 9:**
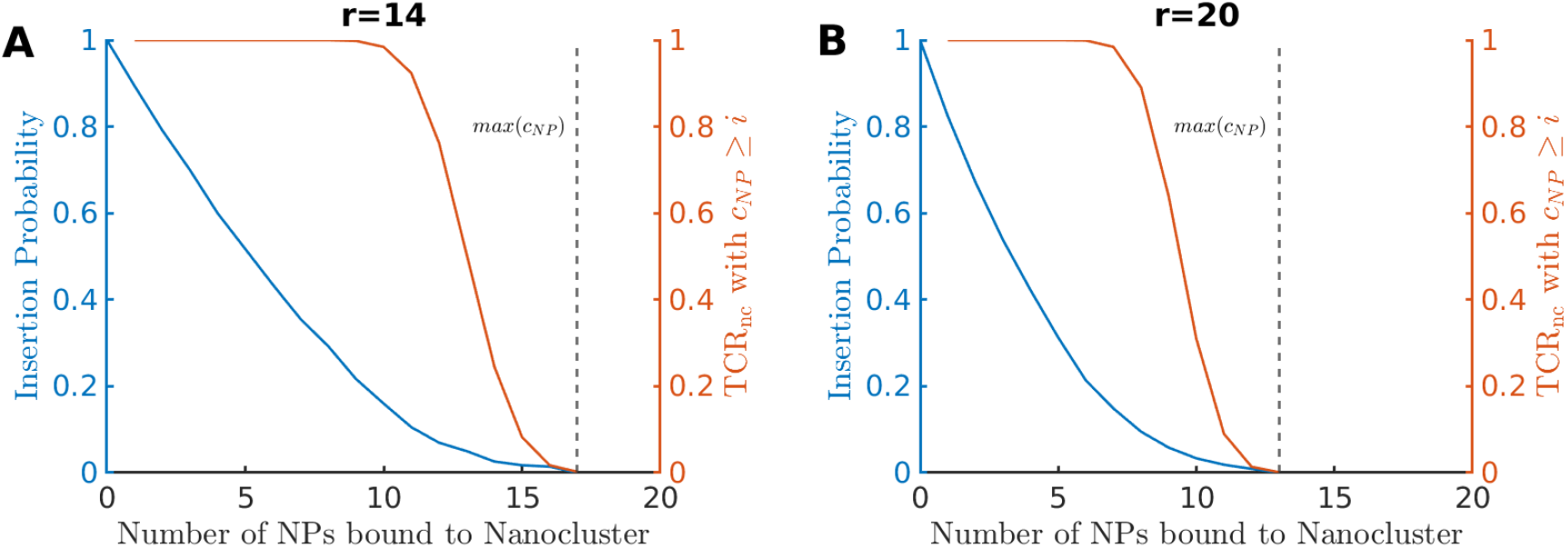
Probabilities associated with NP binding to TCR_nc_. Insertion probabilities associated with TCR_nc_ (blue) and the proportion, *q*(*j*), of TCR_nc_ that are able to bind *j* NPs (red) for (A) *r* = 14 and (B) *r* = 20 nm NPs. Dashed vertical lines represent the theoretical limit of maximum carrying capacity: *M*. In contrast to CAM, the insertion probabilities here denote the proportion of remaining TCRs on the TCR_nc_ following a NP binding event that are available for subsequent binding.

Using the same computations, we also quantified the number of covered TCRs per TCR_nc_ when NP radius is set to *r* = 14 and *r* = 20 nm. These computations reveal that, for *r* = 20 nm, the first NPs to bind to a TCR_nc_ will cover more TCRs than those that bind later (Fig. 10). This is not the case for *r* = 14 nm NPs (results not shown), because *d_TCR_* = 10 nm and the geometry allows only for one single TCR to be contained within the contact area.

**Figure 10:**
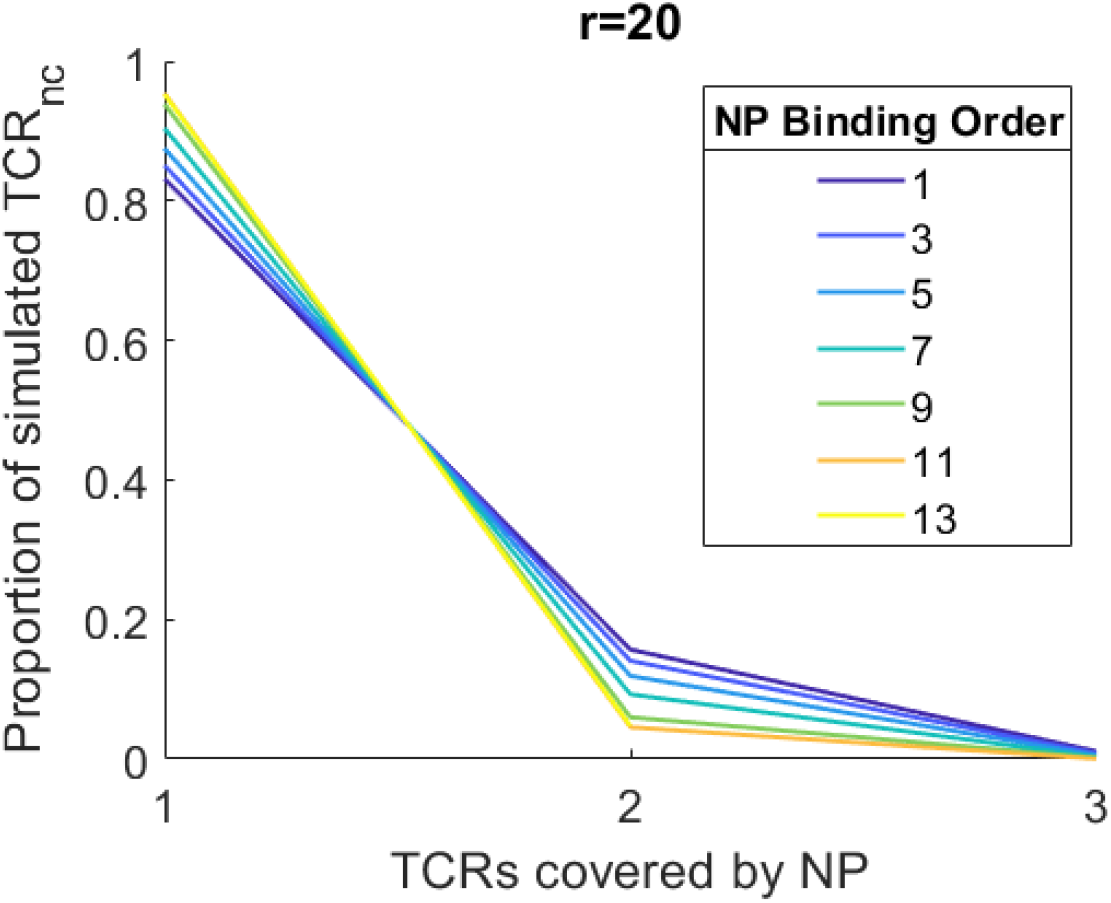
Probability distribution of covered TCRs for every single NP bound to a TCR_nc_. The number of covered TCRs per *r* = 20 nm NPs for each binding event when *d_TCR_* = 10 nm. Each curve is color-coded according to the order of binding specified in the legend.

Notice that there is a very high probability for *r* = 20 nm NPs to cover few TCRs (Fig. 10), a direct consequence of the minimum distance *d_TCR_* = 10 nm imposed between TCRs. Decreasing *d_TCR_* to 5 nm does not change this particular feature of NP binding to TCR_nc_ (Fig. S10). However, the distributions of covered TCRs, in this case, become more distinct between NP bindings for *r* = 14 nm and appear more left-shifted compared to those for *r* = 20 nm NPs. This is to be expected since increasing the size of NPs increases the probability of covering (and thus binding) multiple TCRs, thereby prolonging the dwell time of the NP in the TCR_nc_ and increasing the number of engaged TCRs per NP. This, of course, is done at the expense of TCR_nc_ carrying capacity *c_NP_*: the larger the NPs are, the smaller the *c_NP_* will be. A trade off between these two factors is at a play, but determining how they collectively impact dynamics is investigated at the T cell level.

#### 3.2.3 Continuation Method for Computing the Steady State Distributions of *Y_j_* in the TNM

Deciphering the kinetics of T cell activation requires quantifying the steady states *Ŷ_j_* (*j* = 0,…, *M*) of *Y_j_*, whose dynamics are described by the nonlinear multi-state model given by Eqs. (43)–(45). To do so, we employ here the parameter continuation method [42] and use the vector form

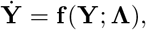

to express the multi-state model, where **Y** = (*Y*_0_,…, *Y_M_*)^*T*^ and **Λ**:= (*K_D_*, *v*, *D*). Solving for the steady states of this system for a given set of parameters is equivalent to solving **f**(**Y**; **Λ**) = **0**, a root-finding problem that is done numerically using Newton’s method [42]. In this analysis, we consider discrete (*D*, *v*) pairs separately, and run their corresponding parameter continuations independently, restricting ourselves to a single parameter: the dissociation constant *K_D_* = *k_off_*/*k_on_*. This parameter essentially determines the distribution of nanoclusters across the *Y_j_* states and, by extension, determines the dynamics of TNM.

Implementing this parameter continuation method for *K_D_* ∈ (10^-2^, 10^4^), when *r* = 20 nm and *v* = 9 (Fig. 11A) vs 13 (Fig. 11B), we obtain the TNM steady state distributions *Ŷ_i_*, for *j* = 0,…, *M*, where *M* = 13 is the maximum carrying capacity of the TCR_nc_. As expected, for higher values of *K_D_*, generated by increasing *k_off_* relative to *k_on_*, an increase in the proportion of low-occupancy steady states *Ŷ_j_* with low *j* indices is observed. Conversely, lowering *K_D_* allows the TCR_nc_ to reach more elevated occupancy steady states *Ŷ_j_* possessing high *j* indices.

**Figure 11:**
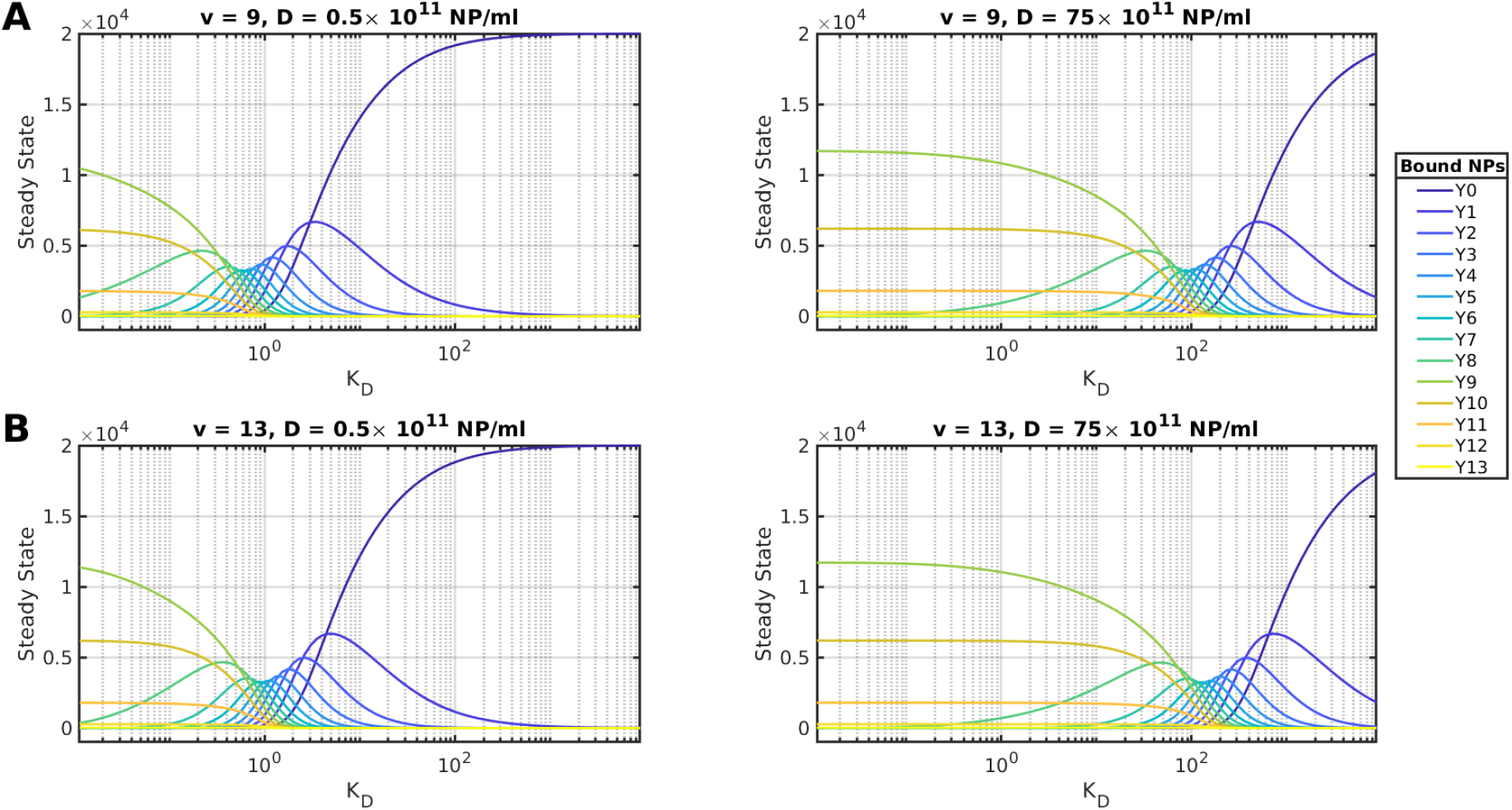
Steady state distributions of the proportion of TCR_nc_ with *j* bound NPs. Plots of the steady state distributions *Ŷ_j_*, *j* = 1,…, *M*, of the TNM computed for NPs with radius *r* = 20 nm and valences (A) *v* = 9, and (B) *v* = 13. Two NP concentrations *D* are used: *D* = 0.5 × 10^11^ NP/ml (left), and *D* = 75 × 10^11^ NP/ml (right). Each curve represents the proportion of TCR_nc_ in each steady state *Ŷ_j_* for a given value of *K_D_*. The parameter continuation method is used to iterate the steady state distributions through a range of *K_D_* values. Each curve is color-coded according to the legend.

By increasing the valence of NPs from *v* = 9 to *v* = 13, while keeping the concentration fixed, to evaluate how that would affect the profiles of *Ŷ_j_*, a moderate shift in the steady state distributions toward higher occupancy states *Ŷ_j_* is generated for given *K_D_* values (Fig. 11). Similarly, increasing *D* while keeping the valence fixed yields a horizontal translation of the steady state distributions to the right, indicating a general shift toward a high occupancy states *Ŷ_j_*.

These outcomes remain roughly the same for *r* = 14 nm NPs (Fig. S11), except for two major differences: (i) the maximum carrying capacity reached between the two radii is not the same (*M*_*r*=20_ = 13 vs *M*_*r*=44_ = 17), implying that *M* is dependent on NP radius, as well as (ii) the moderate change observed in *Ŷ_j_* produced by increasing the valence from *v* = 9 to *v* = 13 for *r* = 20 nm NPs is virtually nonexistent for *r* = 14 nm NPs when valence is increased by relatively similar magnitude from *v* = 8 to *v* = 11 while keeping *D* fixed. These differences thus suggest that there could be a trade-off between size and maximum carrying capacity, that the role of NP valence is limited to CAM for mostly small NPs, and that NP size can enhance the effect of the valence on NP binding to TCR_nc_ in the TNM.

Further examination of these distributions for *r* = 14 and *r* = 20 nm (Figs. S11 and 11, respectively) also reveals that, for smallest values of *K_D_*, some steady states *Ŷ_j_* plateau at different levels, implying that some TCR_nc_ can reach their maximum carrying capacity *M*. For intermediate values of *K_D_*, on the other hand, there are both TCR_nc_ with few as well as many NPs bound, indicating that binding is strong enough to allow for binding of NPs, but not strong enough to reach *M*. Finally, for high values of *K_D_*, most TCR_nc_ end up binding very few NPs.

### 3.3 Outcomes of T Cell Activation Model (TAM)

#### 3.3.1 Estimating Distribution of Bound TCRs per T Cell

We now turn our attention to estimating the number of bound TCRs per T cell for the T cell activation model (TAM), given the number of bound TCR per TCR_nc_. By taking each TCR_nc_ to be independent, it suffices to consider only the distribution of bound TCRs per TCR_nc_ to estimate the global distribution of bound TCRs across the whole cell. Our approach will utilize the distributions of covered TCRs per NP derived from the TNM combined with the CAM estimates of bound TCRs per NP as a function of covered TCRs. This will be done while keeping in mind that the distribution of covered TCRs per NP is dependent on the order of binding of the NP.

To tackle this problem, we let

- *d*(*n*) be the number of bound TCRs per NP as a function of covered TCRs *n*,
- 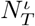 be the number of covered TCRs by the *ι*^th^ NP,
- 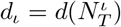 be the number of TCRs bound by the *ι*^th^ NP,
- *D_j_* be the number of bound TCRs per TCR_nc_ with *j* NPs attached,
- 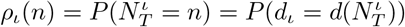 be the discrete probability distribution of covered TCRs per NP, and
- ʔ be the total number of bound TCRs per T cell.

These quantities can be approximated by using an averaging approach. This can be done by employing the TNM and CAM to compute estimates of these quantities using the following parameters

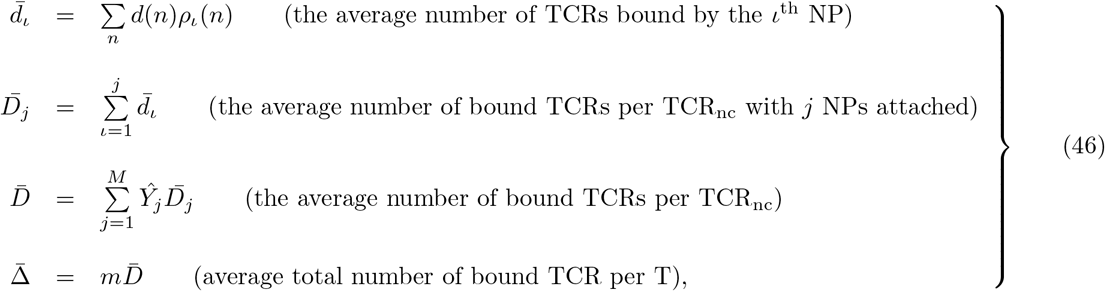

where *m* is the total number of TCR_nc_ and 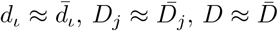 and 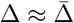.

Even though this approach represents a straightforward method for estimating Δ, considering the distribution of this variable instead will likely capture a variety of responses across a T cell population, making the model more reliable. Obtaining a distribution for Δ, however, is complicated by the fact that our initial distributions of covered TCRs *ρ_ι_* is discrete, whereas the function *d*(*n*) that scales *ρ_ι_* to obtain the distributions of bound TCRs is computed from CAM and thus may yield non-integer values. We resolve this issue by fitting a continuous distribution *f_ι_* to the numerical estimates of the discrete distribution of *d_ι_*, followed by using *f_ι_* to compute the probabilities of bound TCRs at integer values. For the choice of fit function *f_ι_*, we select the cumulative distribution *F_ι_*(*x*) of the two-parameter gamma distribution, given by

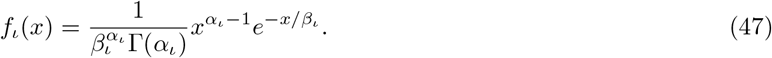

The fitting of *F_ι_* to the discrete cumulative distribution associated with *d_ι_* is then done in two steps. In the first step, we allow independent estimates of *α_ι_* and *β_ι_* for each bound NP (*ι* = 1,…, *c_NP_*); this produces *β_ι_* values that are very similar to each other (results not shown). Motivated by the closeness of *β_ι_* values, in the second step, we calculate the average

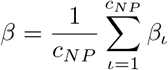

and use it to fit (Fig. 12) *F_ι_* (orange curves) to the discrete cumulative distribution associated with *d_ι_* (blue squares) for each bound NPs and obtain new estimates of *α_ι_* (*ι* = 1,…, *c_NP_*). Doing so produces fits between continuous and discrete cumulative distributions that match really closely, independent of the order of NP binding.

**Figure 12:**
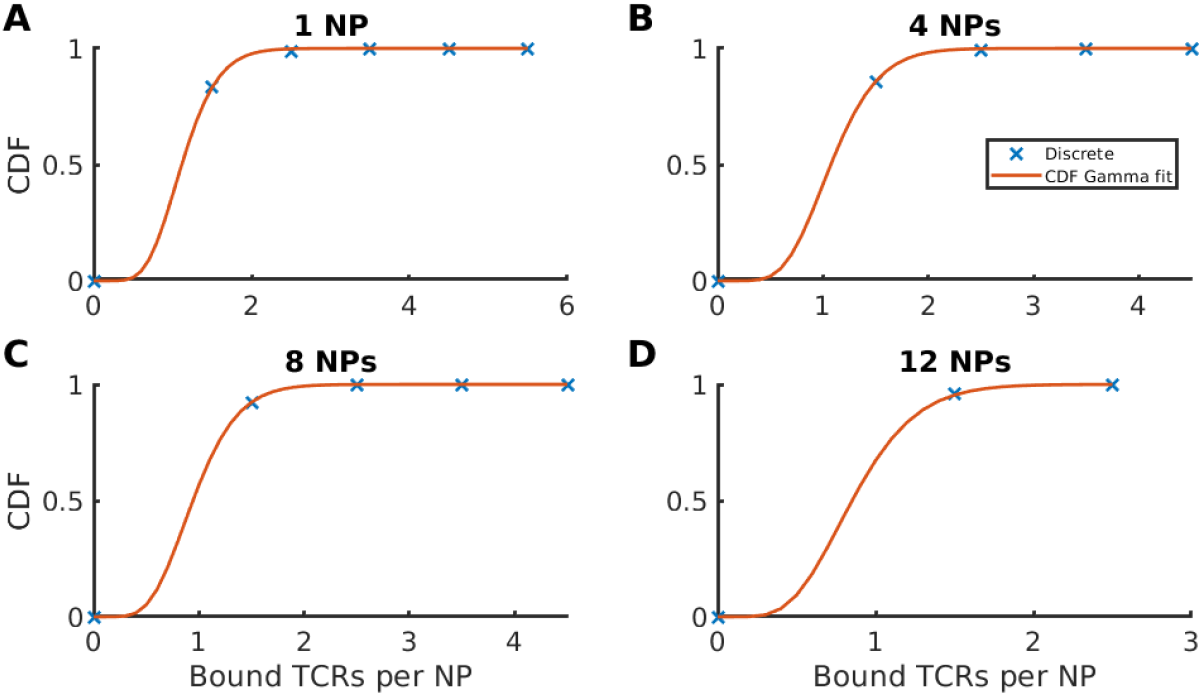
Cumulative distribution function for the discrete probability of bound TCRs per NP fit to a continuous cumulative gamma distribution function. The distribution varies for each NP based on the order of binding to a TCR_nc_, with each biding producing a new combination of (*α_ι_*, *β_ι_*), *ι* = 1,…, *c_NP_*. Because the change in *β_ι_* is small between bound NPs, we instead use the average *β* to fit the cumulative gamma distribution (orange curves) to the discrete cumulative distribution of covered TCRs defined by *d_ι_* (blue squares) for all NPs (*ι* = 1,…, *c_NP_*). Plots shown correspond to the distributions of the (A) first, (B) fourth, (C) eighth and (D) twelfth NPs that bind to a TCR_nc_.

The choice of the gamma distribution is motivated by the fact that the distribution of *D_j_* can be efficiently calculated via convolutions of the independent distributions of *d_ι_* for *ι* = 1,…, *j*. Specifically, if the random variable *d_ι_* of bound TCRs to *ι*^th^ NP is sampled from the distribution

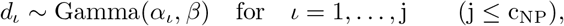

which happens to be the case here (given that *β_ι_* values are similar and the average *β* is chosen), then

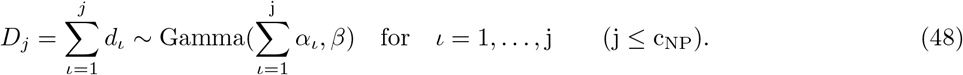

One can thus take advantage of this property of the gamma distribution to compute *D_j_* in a straightforward manner.

To further examine the performance of the continuous gamma distribution, we compare the density of bound TCRs predicted by this distribution and the one calculated using the discrete distribution *d_ι_* at the TCR_nc_ level (Fig. 13). The continuous gamma distribution successfully approximates the discrete distribution *d_ι_* very closely at intermediary levels of bound NPs. When *ι* gets large (compare 1 NP (A) to 12 NP (D) in Fig. 13), the predicted distribution slightly underestimates the actual number of bound TCRs per TCR_nc_. However, this discrepancy is mitigated by the fact that TCR_nc_ are unlikely to bind that many NPs (Fig. 9). It is important to note here that the Gaussian distribution has also been tested, but the gamma distribution out-performed it (results not shown; see [42] for more details).

**Figure 13:**
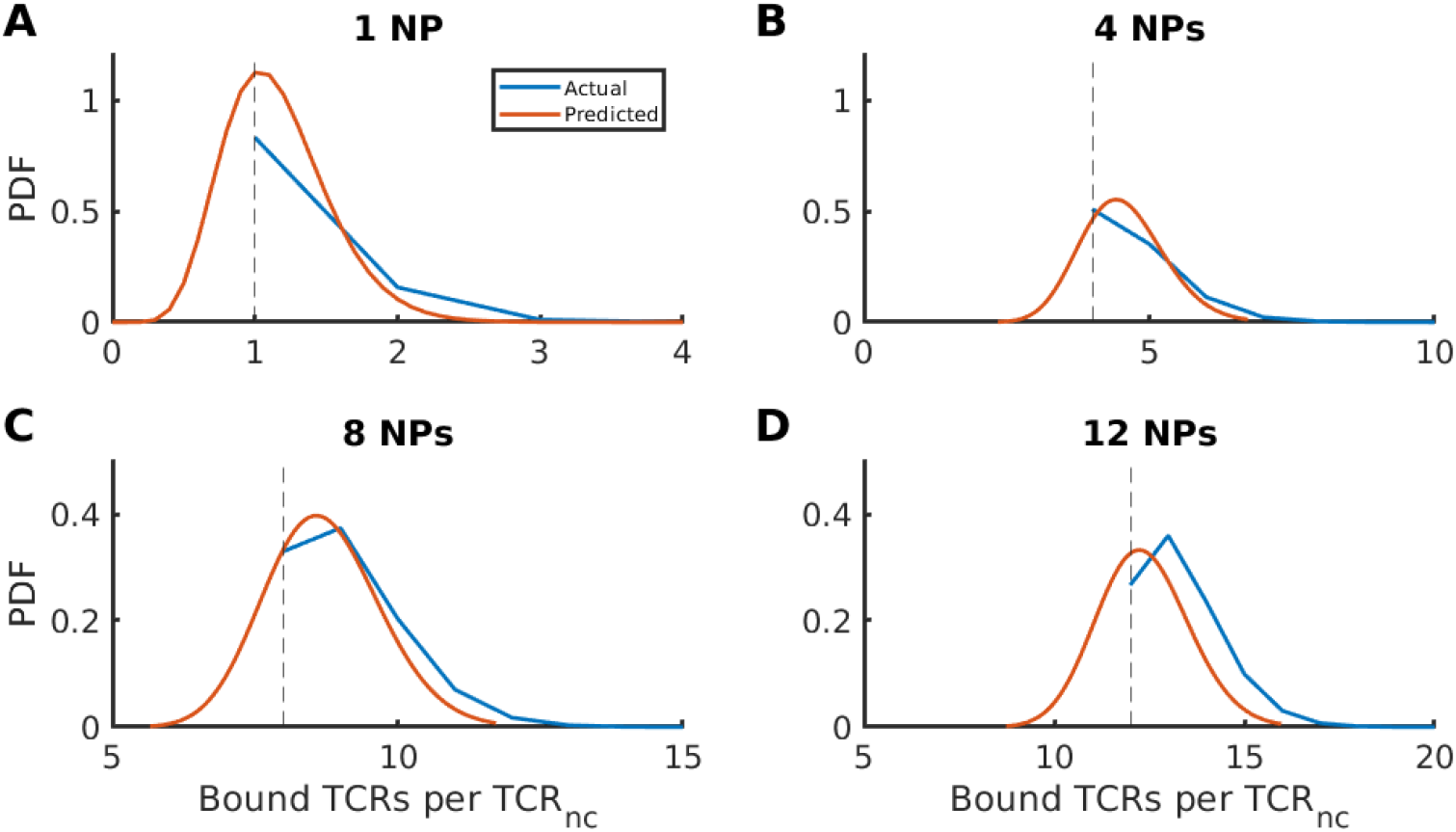
The probability density function of bound TCRs per TCR_nc_. The distribution predicted by the fitted gamma distribution (orange curve) and actual distribution of *d_ι_* (blue) for the special case where all covered TCRs are bound. Distributions vary depending on the number of NPs bound to the TCR_nc_. Shown are the cases for (A) 1 NP, (B) 4 NPs, (C) 8 NPs and (D) 12 NPs bound to the TCR_nc_.

We can now scale this TCR_nc_ distribution to the whole T cell by recognizing that each TCR_nc_ is independent. Recall that for a total of m TCR_nc_ per T cell, *Ŷ_j_* represents the proportion of TCR_nc_ that contain *j* bound NPs at steady state, where *j* ≤ *M*, the maximum carrying capacity of the TCR_nc_. Based on this, we have

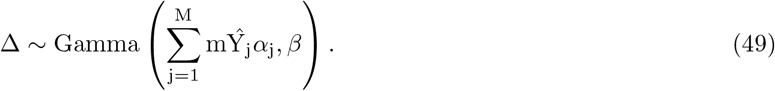

#### 3.3.2 Subthreshold vs Suprathreshold T Cell Activation Determined by TAM

Now that the distribution of number of covered TCRs per T cell Δ can be computed using Eq. (49), we can use it to directly fit the multiscale model to NP-dependent IFN_*γ*_ dose-response curves generated by stimulating 25000 CD8^+^ T cells with different concentrations of NPs of radius *r* = 14 and *r* = 20 nm (Fig. S3). The fitting involves estimating the parameters *μ*, 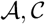, *K_D_* and 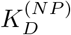, defined by Eqs. (3), (14) and (42). This is done by matching model outcomes to data at all valences simultaneously for each radius using MCMC techniques with 15000 iterations initiated from those average values obtained using Eq. (46). The results obtained show doseresponse curves that resemble those obtained experimentally reasonably well at various valences of both radii except for those at low valences between *v* = 9 and *v* = 13 when *r* = 14 nm, where a jump from subthreshold to suprathreshold responses are observed (Fig. S3; compare dashed and solid curves). Note here that, for *r* = 20 nm NPs, the sub-to-suprathreshold jump between *v* = 13 to *v* = 61 is better captured by the model but the change in valence is significantly higher here compared to that for *r* = 14 nm NPs. The jump in the simulations, however, becomes pronounced for *r* = 14 nm when plotted with respect to pMHC concentration rather than NP concentration (results not shown).

The discrepancy at low valences when *r* = 14 nm could be due to one or a combination of four potential factors: (1) presence of positive cooperativity in the binding of NPs to TCR_nc_, which we have shown before not to be the case; (2) the model is purely based on the geometry of interaction between NPs and T cells and thus does not account for signaling events downstream of TCR triggering that could act in synergy in amplifying the response; (3) the minimum distance between TCRs *d_TCR_* = 10 is too big to allow for polyvalent binding between *r* = 14 nm NPs and TCR_nc_; and (4) the experimentally generated dose-response curves are formed from single data points at each NP concentration for a given NP size, which may cause the sub-to suprathreshold jumps to occur between other valences.

Decreasing the minimal distance between TCRs to *d_TCR_* = 5 nm generates results that are similar to those seen with *d_TCR_* = 10 nm; indeed, allowing *r* = 14 nm NPs to form polyvalent bonds with TCR_nc_ by cutting *d_TCR_* in half does not improve the ability of the model to capture the jump between sub-and suprathreshold responses for *r* = 14 nm. In the next sections, we will examine point (4) and address point (2) in the Discussion.

#### 3.3.3 Parameter Sweeping for Detecting Jumps Between Dose-Response Curves

With the presence of only one data point per concentration, valence and radius, one needs to explore if the jump from subthreshold to suprathreshold responses do indeed occur between valences that could be as low as *v* = 8 and *v* = 11. This can be accomplished by simulating TAM, parametrized by those values obtained from the MCMC fitting, and plotting the heat-maps of T cell IFN*γ* production generated at steady states for both radii *r* = 14 (Fig. 14A) and *r* = 20 (Fig. 14B) while varying both the concentration *D*, and valence *v*. Supper-imposing the level curves on those heat-maps allows us to uncover if prominent changes in IFN*γ* production occurs and, if yes, in what biophysically reasonable parameter regimes. When level curves are close in a given parameter regime, it indicates that there is a drastic change in the release of IFN*γ* in that regime.

**Figure 14:**
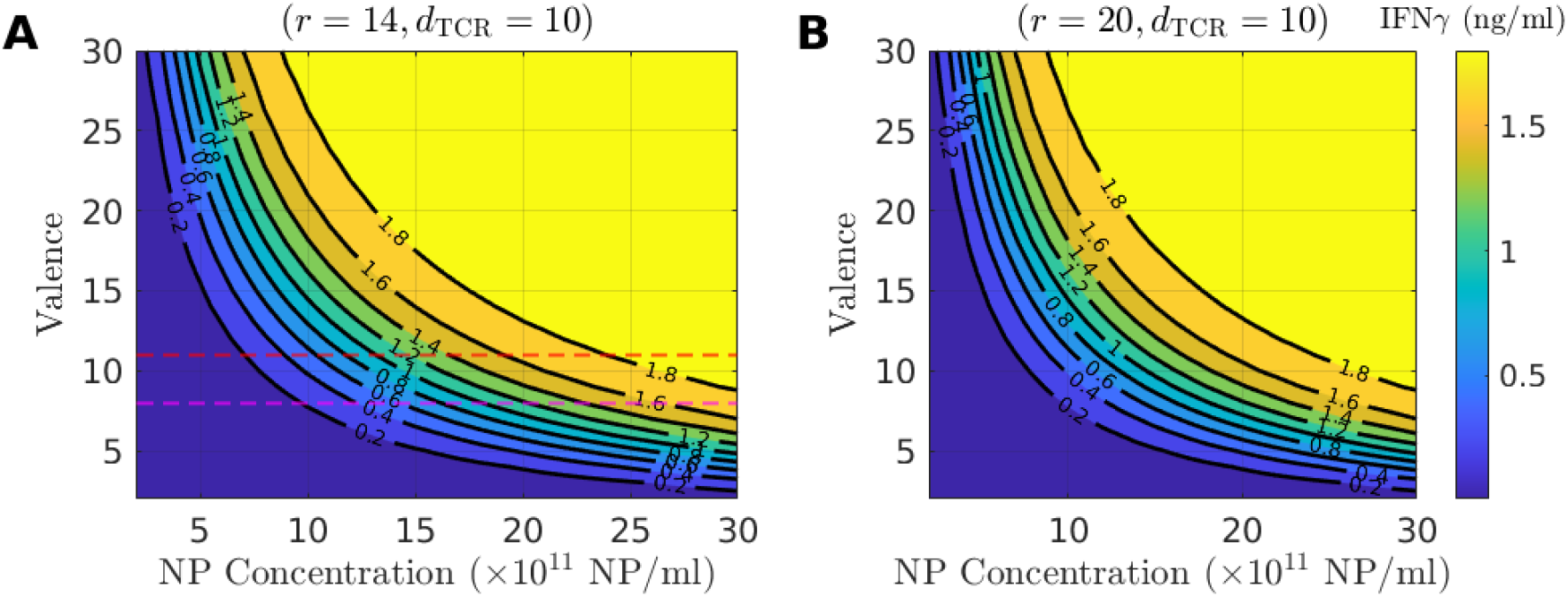
Dependence of T cell IFN*γ* production on NP concentration, valence and size as determined by TAM. Heat-maps of IFN*γ* production when varying NP concentration and valence for (A) *r* = 14 nm NPs, and (B) *r* = 20 nm NPs with *d_TCR_* = 10 nm. Heat-maps are color-coded according to the color-bar in each panel. Black lines are the level curves labeled according to the magnitude of IFN*γ* production (in ng/ml). Horizontal lines in A indicate valences *v* = 8 (pink) and *v* = 11 (red).

By plotting these level curves for both radii (Fig. 14), we find that generating pronounced increases in the level of IFN*γ* over a whole range of NP concentrations for both radii is possible. Indeed, one can obtain such an increase in IFN*γ* between two small valences for *r* = 14 nm NPs (Fig. 14A), as long as the difference in valences is large enough, especially compared to that between *v* = 8 and *v* = 11 (e.g., *v* = 5 vs *v* = 12). Similar results can be obtained with *r* = 20 nm NPs (Fig. 14B), but with less discrepant small valences like *v* = 6 vs *v* = 11. Al in all, these results indicate that, geometrically speaking, TAM can alone generate prominent jumps between IFN*γ* dose-response curves associated with two low valences at a given NP size; these jumps, however, remain not as pronounced as those seen experimentally. One can then postulate that the geometry of NP polyvalent biding to T cells could belong to the proper parameter regime of pMHC density to facilitate T cell activation by working in synergy with downstream signalling pathways to produce IFN*γ* jumps.

### 3.4 Discussion

In this study, we modeled the polyvalent binding of NPs to T cells based almost exclusively on the geometrical properties of this system. Taking advantage of the mathematical framework of polyvalent receptor-ligand interaction presented in [40], we developed a multiscale model of T cell binding to NPs and incorporated the relevant geometries associated with this interaction at each scale. There were a total of 3 different scales that we considered: the contact area level (i.e., the area between the NPs and T cells) which includes the TCR-pMHC binding, the TCR nanocluster level (the nanoscale structures formed by TCRs on the membrane of T cells), and the T cell level. Although the model is adapted to the interactions between T cells and NPs, the framework of the model is generalizable to any polyvalent binding system possessing similar geometries at these three different spatial scales.

The model was affinity-centric, linking the activation of a single T cell to the dissociation constant *K_D_* of the TCR-pMHC complex and the total number of bound TCRs on T cells. With this multiscale model, we combined both analytical and computational techniques to quantify several aspects of NP interactions with T cells at each scale. That included quantifying the insertion probabilities of pMHCs and NPs in the CAM and TNM models, respectively, determining the dwell-time NPs spent bound to TCR_nc_ as a function of dissociation constant, as well as computing the distributions of covered and bound TCRs per NP for two NP sizes: *r* = 14 and *r* = 20 nm. The latter required fitting the continuous gamma distribution to the non-integer discrete distribution of bound TCRs.

Combining CAM with TNM dynamics, we went on to explore how the model performs when compared to dose-response curves of IFN*γ* release from T cells upon NP stimulation. Embedding TCR kinetic proofreading (Fig. S1) along with TCR triggering (Fig. S2) into the multiscale model revealed that at those valences used experimentally (e.g., *v* = 8 and *v* = 11 for *r* = 14 nm NPs), jumps from subthreshold to suprathreshold responses over a whole range of NP concentrations were not possible to capture properly with the model when NP radius is small (Fig. S3). Different MCMC strategies were used to estimate model parameters but without yielding any improvement to the quality of the fits. We then verified that the angle (45°) of the contact area was optimized for producing a large difference in effective valence between *v* = 8 and *v* = 11 (Fig. 7). Our analysis of cooperativity also demonstrated that for low valence, cooperativity tended to initially decrease with respect to NP-valence.

Because experimentally measured IFN*γ* dose-response curves (Fig. S3) were formed by using one single data point per NP concentration for each NP valence and size, we hypothesized that the estimates of pMHC-density dependent thresholds may not be precise. To explore this, we plotted the heat maps of IFN*γ* responses over a whole range of NP valences and NP concentrations for both radii *r* = 14 and *r* = 20 nm using the multiscale model (Fig. 14). Our results highlighted that jumps in IFN*γ* responses can occur across small valence differences. Although these jumps were not as pronounced as those associated with subthreshold to suprathreshold pMHC densities [23], they could potentially be sufficient to facilitate T cell response downstream. We hypothesized that the binding kinetics of NPs under the geometrical constraints considered in this study may act in synergy with downstream subcellular effectors, such as Zap70 (not included in the model) to activate T cells. Evidence suggests that the gene regulatory network underlying the dynamics of Zap70 phosphorylation and activation possesses a hysteresis in the form of a bistable switch [46,47]. The moderate jumps induced by the multiscale model could be necessary to push Zap70 across the activation threshold (saddle-node bifurcation) in the bistable switch and thus induce this apparent pMHC-density dependent threshold seen experimentally.

The TNM developed here assumed that the TCR_nc_ exist prior to pMHC-NP binding. Evidence for the existence of these nanoclusters has been previously suggested [33]. In our analysis, T cells are assumed to be autoantigen-experienced, implying that the TCR_nc_ are fully formed, allowing us to quantify this polyvalent binding between NPs and T cells including cooperativity. Interestingly, it has been also shown, that upon encountering pMHCs, these nanoscale arrangements of TCRs on the T cell grow in size [30]. This process is believed to be related to the formation of the immune synapse. This increased level of aggregation is thought to enhance the efficacy of interactions between T cells and antigen presenting cells, both by providing a stimulating environment for further binding events to occur and enhancing the signaling of TCR within the aggregate [30]. To consider the effects of nano- and microclustering on binding and triggering it would be fundamentally important to investigate dynamically how these clusters change in time upon activation, especially upon NP stimulation. Estimating the distribution of the number of TCRs per TCR_nc_, before and after T cell stimulation, one can then determine how the distribution changes upon activation and subsequently evaluate the relation between TCR triggering and TCR microclustering. It is important to point out that although only the qualitative changes in the distribution can be considered, the results in [30] were merely qualitative, as the technique there used to determine the sizes of the TCR_nc_ was shown to generally underestimate the number of TCRs in a nano- or microcluster. Nonetheless, such analysis will permit further quantification of these processes (called agglomerations), and unravel its contribution to T cell activation.

While the multiscale model considered here is affinity-centric, it has been suggested that in the context of an immune synapse, pMHCs can serially engage 200 TCRs [5,9]. While serial engagement may be specific to immune synapse, it is very plausible that this mechanism of interaction also occurs between T cells and the pMHC-coated NPs in a polyvalent manner. Modifying the multiscale model to include this aspect and comparing the outcomes of this new model and how it performs against data with the one presented here represents an interesting new avenue that one can also explore to see if serial engagement can contribute more to the pMHC-density dependent thresholds that produce the jumps between the IFN*γ* dose-response curves than the affinity-centric model or not.

This study was quantitative in nature; it allowed us to develop a theory that helped decipher how nanoparticles can polyvalently bind to TCRnc and induce T cell activation. Although, it was only limited to the geometry of the interaction, it was still able to elucidate key aspects of this system and highlight the implications of different factors involved.

## Acknowledgment

This work was supported by the Natural Sciences and Engineering Council of Canada (http://www.nserc-crsng.gc.ca/index_eng.asp) discovery grant to AK. The authors would like to thank Dr. Pere Santamaria (Department of Microbiology Immunology and Infectious Diseases, University of Calgary) and his laboratory members for providing us with the IFN*γ* response data.

