## Supplemental Figures for "Theoretical Quantification of the Polyvalent Binding of Nanoparticles Coated with Peptide-MHC to TCR-Nanoclusters"

March 2022

### Symbols and Definitions

| **Symbol** | **Definition** |
|---|---|
| CAM | Contact Area Model. |
| TNM | T-cell Nanocluster Model |
| TAM | T-cell Activation Model |
| NP | Nanoparticle. |
| TCR | T cell Receptor. |
| $\text{TCR}_{NC}$ | TCR Nanocluster. |
| $v$ | Valence. |
| $\tilde{v}$ | Effective valence (number of pMHC in contact area). |
| $d_{TCR}$ | Minimum separation between TCRs. |
| $n_{TCR}$ | Number of TCRs per $\text{TCR}_{\text{NC}}$. |
| $X_i$ | CAM Markov states. Proportion of contact areas containing $i$ bound TCR-pMHC complexes. |
| $N_T$ | Number of TCRs in the contact area. |
| $N = \min(N_T, \hat{v})$ | Maximum number of possible pMHC-TCR complexes for a given contact area |
| $P_i$ | Insertion probabilities of the CAM. Represents the proportion of pMHC in the contact area that are available for binding after $i$ pMHC-TCR complexes have already been formed |
| $Y_i$ | Nanocluster model Markov states. Proportion of $TCR_{NC}$ with $i$ bound NPs. |
| $c_{NP}$ | NP carrying capacity of a $\text{TCR}_{\text{nc}}$ (maximum number of NPs that can bind to a given $\text{TCR}_{\text{nc}}$) |
| $M = \max(c_{NP})$ | Theoretical maximum carrying capacity for *any* $\text{TCR}_{\text{nc}}$ |
| $d(n)$ | Number of bound TCRs pe NP as a function of covered TCRs $n$. |
| $N_T^\iota$ | Number of covered TCRs by the $\iota^{\text{th}}$ NP. |
| $d_\iota = d(N_T^\iota)$ | Number of bound TCRs by the $\iota^{\text{th}}$ NP. |
| $D_j$ | Number of bound TCRs per $\text{TCR}_{\text{nc}}$ with $j$ NPs attached. |
| $\rho_\iota(n)$ | Discrete probability distribution of covered TCRs per NP. |
| $\Delta$ | Total number of bound TCRs per T-cell. |

Table S1: Table of important symbols and definitions.

### Additional Figures

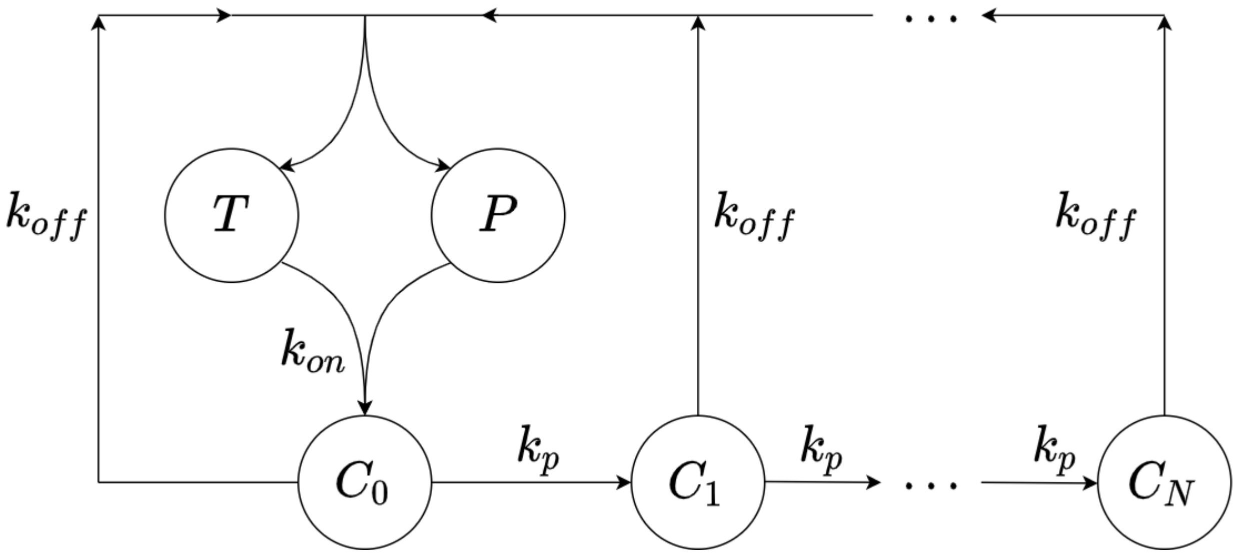


Figure S1: Kinetic proofreading Markov model. Free pMHC ($P$) and TCR ($T$) bind at a rate $k_{on}$ to form a complex ($C_0$). This complex can undergo phosphorylation at a rate $k_p$ and become triggered once $N$ phosphorylation events have occurred. TCR-pMHC complexes need to be triggered in order to contribute to T cell activation and IFN$\gamma$ production. At any point, the complex can unbind at a rate $k_{off}$.

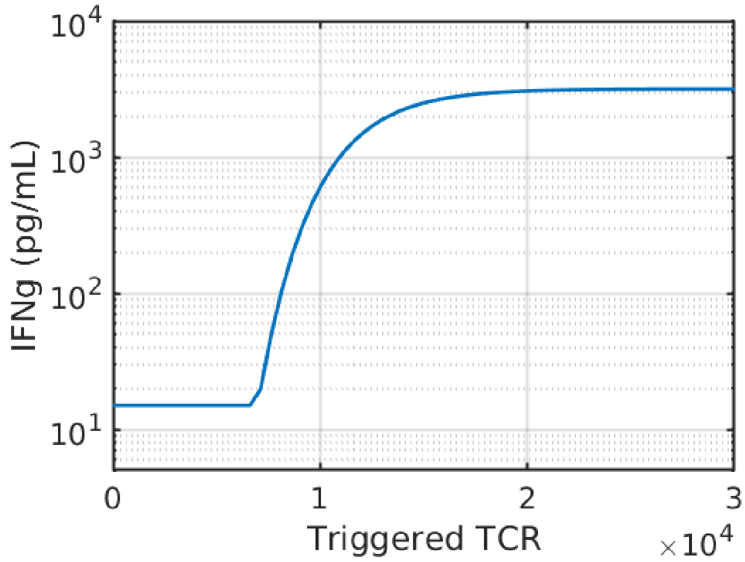


Figure S2: IFN$\gamma$ production as a function of triggered TCRs as defined by Eq. (3).

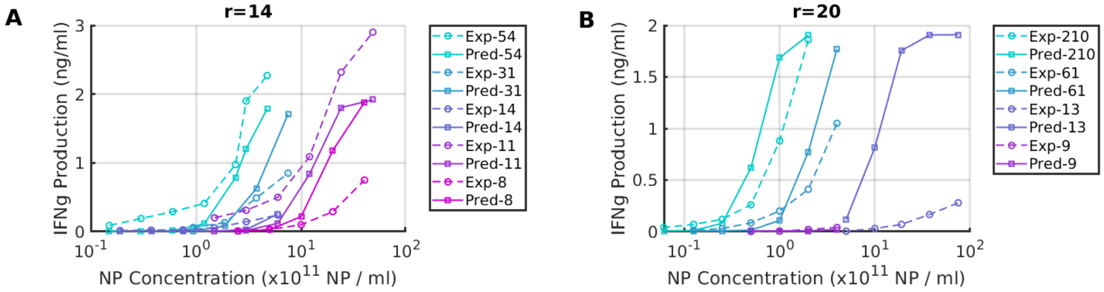


Figure S3: IFN$\gamma$ dose-response curves and model predictions for $d_{TCR} = 10$ nm. Model parameters are estimated by fitting the multiscale model (solid curves) to IFN$\gamma$ dose-response curves (dashed curves) when (A) $r = 14$ (excluding $v = 14$), and (B) $r = 20$ nm. Each curve corresponds to a specific valence $v$ and color-coded according to the legend of each panel. For parameter distributions, see Fig.S12. All experimental data are derived from [23].

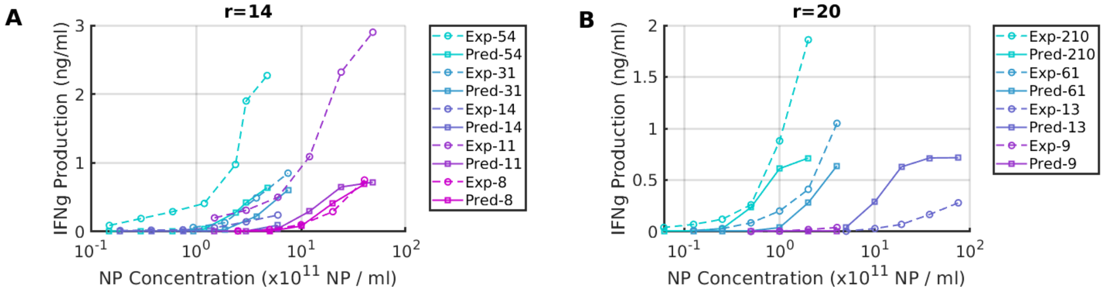


Figure S4: The same as in Fig. S3 except that $d_{TCR} = 5$ nm. All experimental data are derived from [23].

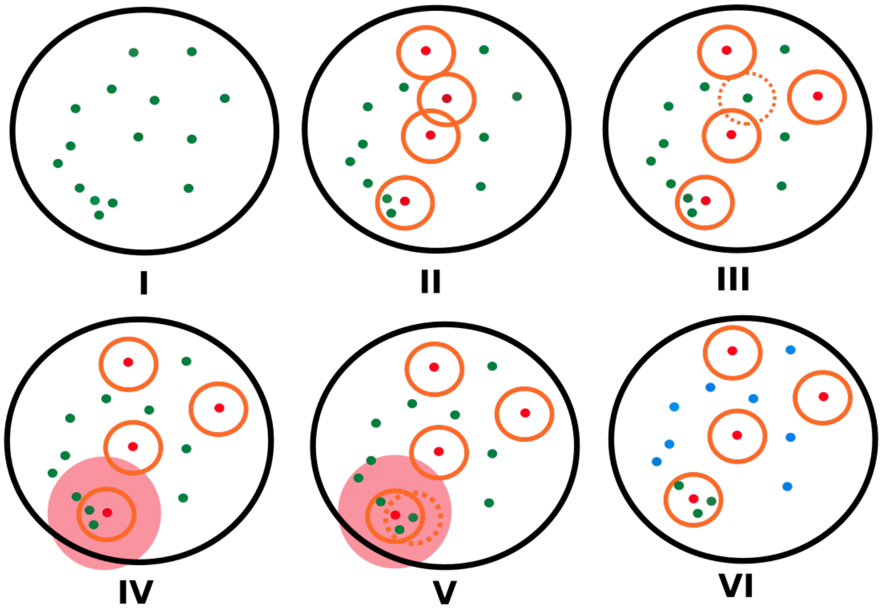


Figure S5: Illustration of Monte Carlo simulations for estimating contact area insertion probabilities. (I) A random configuration of pMHCs within the contact area (green dots). (II) Random configuration of bound TCRs (orange circles). (III) The re-positioning of any overlapping TCRs (dotted circles). (IV-V) TCR positions evolve to the most crowded regions defined by the neighborhood of each TCR (shaded regions). (VI) Counting the number of pMHCs that are neither bound, nor covered by TCRs (blue dots).

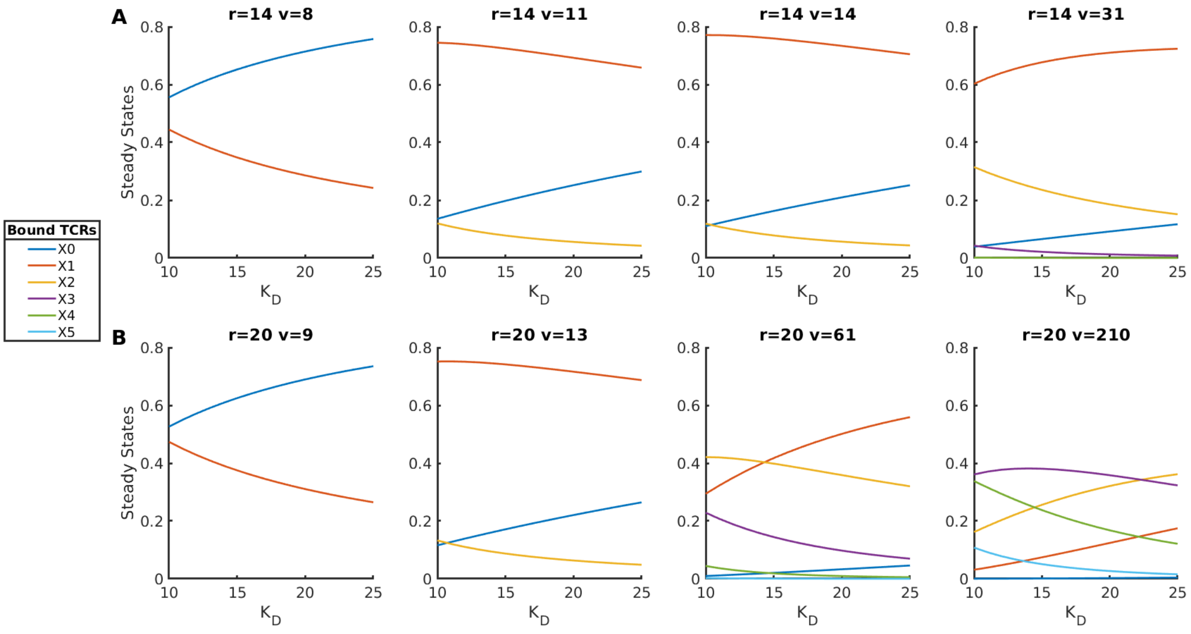


Figure S6: Steady states $\hat{X}_i$, $i = 1, \ldots, N$, of CAM for $d_{TCR} = 5$ nm. Results are displayed in a manner similar to that shown in Fig. 3.

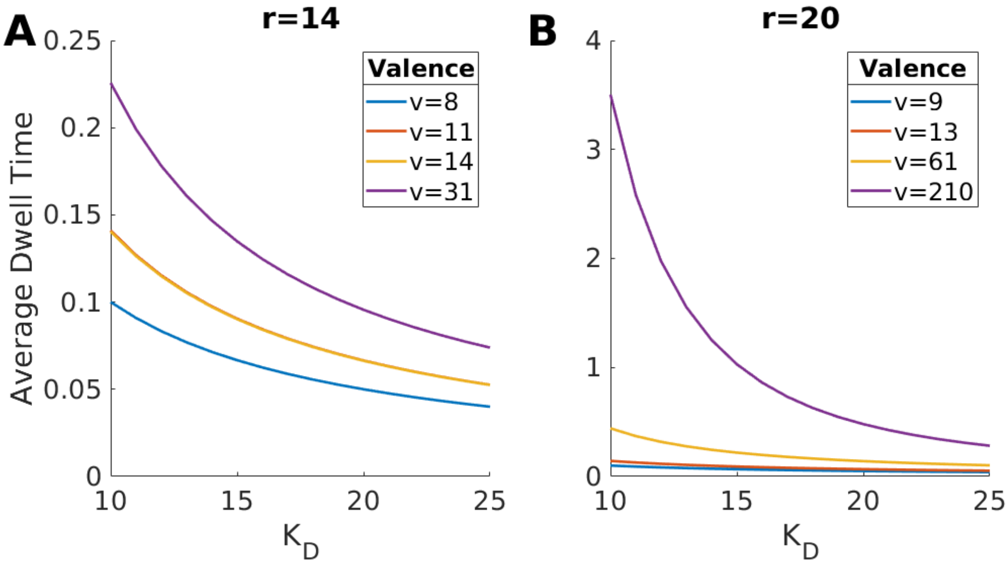


Figure S7: Average dwell times of NPs within the contact area $\hat{\tau}$ for $d_{TCR} = 5$ nm. As in Fig. 5, curves are color-coded according to the legend in each panel.

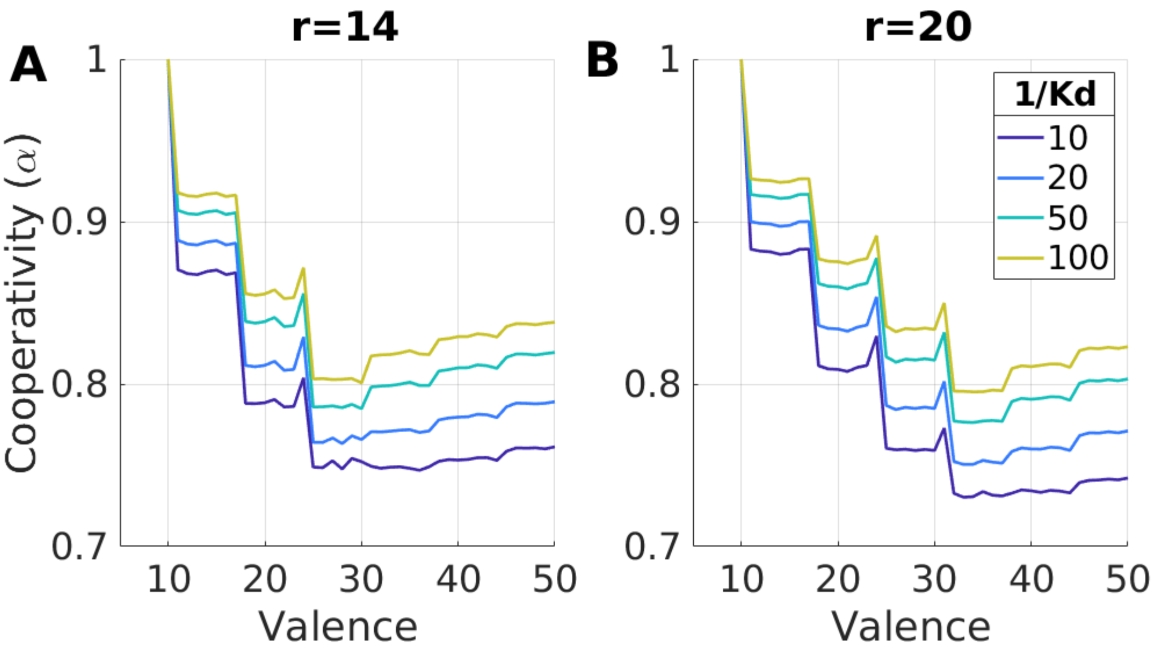


Figure S8: The average cooperativity ($\overline{\alpha}$) computed for $d_{TCR} = 5$ nm and plotted as a function of NP-valence for various values of $K_D$ when (A) $r = 14$ and (B) $r = 20$ nm.

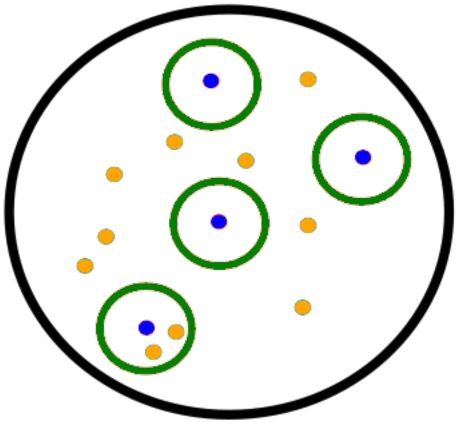

Figure S9: The $\text{TCR}_{\text{nc}}$ geometry is similar to the CAM geometry. We define a radius of $r_{\text{nc}} = 200$ nm (black circle) containing 20 TCRs (yellow: unbound; blue: bound) to which the NPs (green circle) can bind to. We calculate insertion probabilities using the same method outlined in Fig. S5.

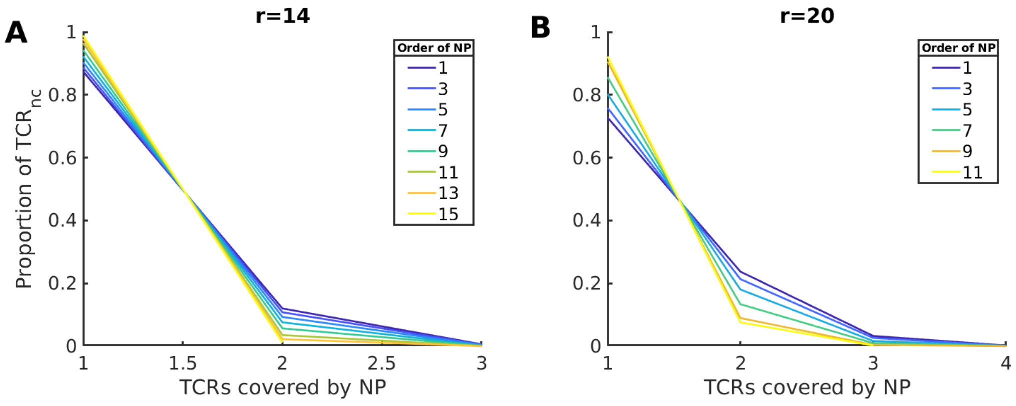


Figure S10: Distribution of covered TCRs for (A) $r = 14$, and (B) $r = 20$ nm NPs on $\text{TCR}_{\text{nc}}$, where $d_{TCR} = 5$ nm.

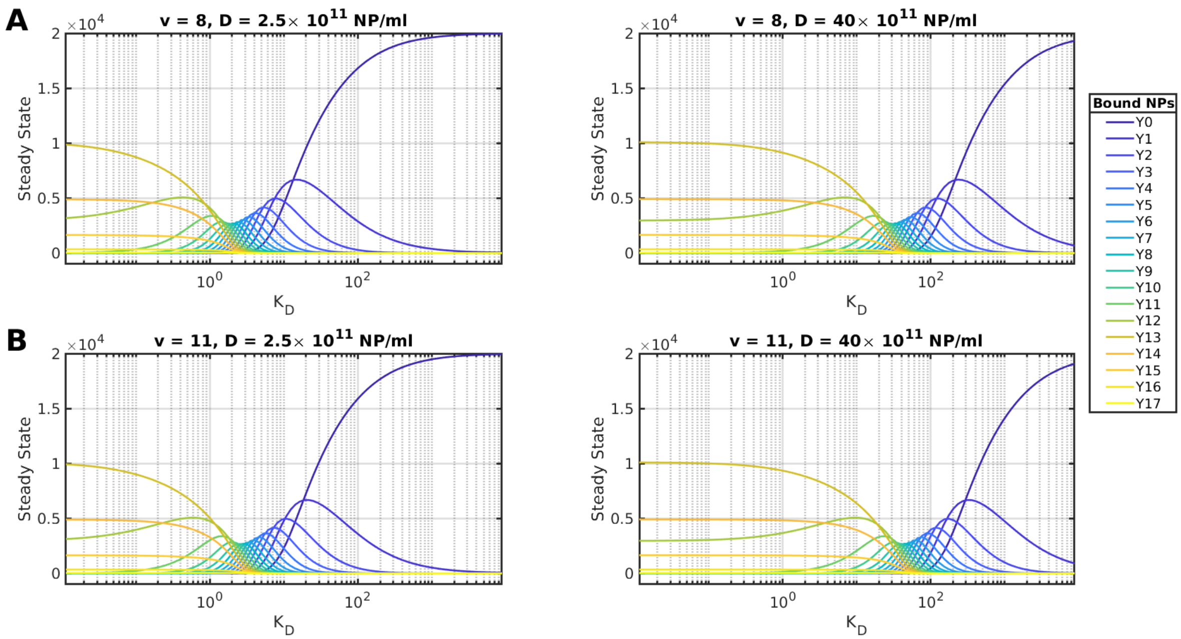


Figure S11: Steady state distributions of the number of NCs with $j$ bound NPs. Plots of the steady state distributions $\hat{Y}_j$, $j = 1, \ldots, M$, of the TNM computed for NPs with radius $r = 14$ nm and valences (A) $v = 8$, and (B) $v = 11$. Two NP concentrations $D$ are used: $D = 2.5 \times 10^{11}$ NP/ml (left), and $D = 40 \times 10^{11}$ NP/ml (right). Each curve is color-coded according to the legend.

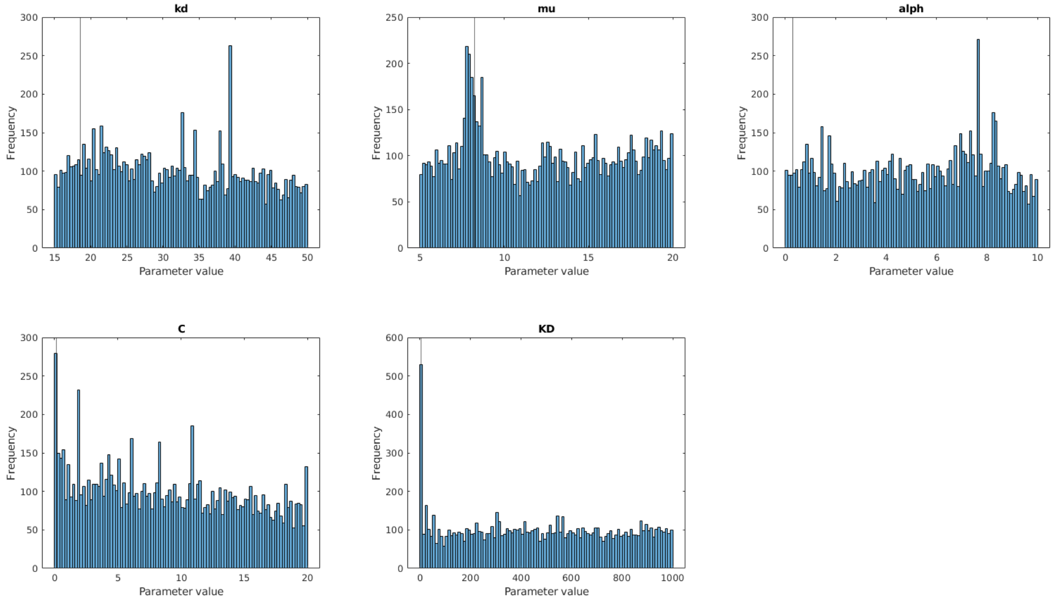


Figure S12: Distribution of sampled parameters from MCMC estimates for $r = 14$ nm fitting. Results of the fit are shown in Fig. S3

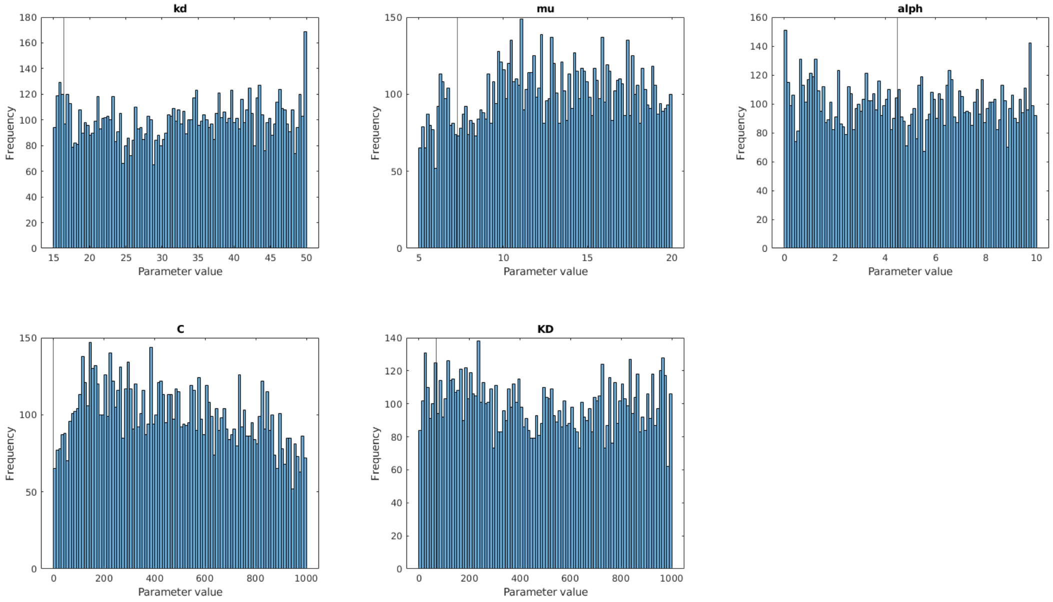


Figure S13: Distribution of sampled parameters from MCMC estimates for $r = 20$ nm fitting. Results of the fit are shown in Fig. S4
